# Syntaxin-2 coordinates endolysosomal trafficking to balance phagocytic uptake and phagolysosomal clearance in macrophages

**DOI:** 10.1101/2024.06.08.598083

**Authors:** Suman Samanta, Abhrajyoti Nandi, Rupak Datta, Subhankar Dolai

## Abstract

Phagocytosis engulfs receptor-bound particles by structuring phagocytic cups that expand to form phagosome vesicles. Phagosomes fuse with endosomes and lysosomes to gradually mature into acidic, hydrolase-enriched phagolysosomes for content degradation. While an essential cellular process for eliminating unwanted substances to secure host defense and organismal homeostasis, defective or uncontrolled phagocytosis can be detrimental. Here, we report syntaxin-2 (Stx2), a poorly characterized SNARE in phagocytes, defines the course of macrophage phagocytosis by coordinating cell surface receptor density, phagocytic cup expansion, and phagosome maturation. Stx2 is expressed on the plasma membrane, phagosomes, early endosomes, and to some extent on recycling endosomes, late endosomes and VAMP4-positive compartments. Stx2 knockdown (Stx2-KD) augments engagement and uptake of IgG-opsonized particles by enhanced surface expression of Fc receptors (FcR) and efficient expansion of phagocytic cups. These events are driven by increased recycling of FcR, and enhanced delivery of early endosomes and VAMP4-positive post-Golgi compartments to phagocytic cups. Interestingly, Stx2-KD macrophages exhibit reduced secretion of pro-cathepsins with concomitant increase in lysosome content. However, Stx2 depletion prevents phagosome coalescence with late endosomes and lysosomes to impair maturation by depleted acquisition of cathepsins and reduced acidification. Consequently, macrophages depleted of Stx2 manifest uncontrolled uptake of IgG-opsonized *Escherichia coli* and impaired digestion resulting in increased bacterial load. Collectively, we uncovered a previously unknown role of Stx2 as a critical balancer of phagocytic uptake and phagolysosomal clearance in macrophages, suggesting that it could be an attractive target for modulation of phagocytosis plasticity and to control aberrant phagocytosis.

## INTRODUCTION

Phagocytosis, a conserved cellular process, engulfs and eliminates large (>0.5 µm in diameter) extracellular particles^1,2^. While various epithelial and non-epithelial cells display phagocytic activity, professional phagocytes of myeloid origin, such as macrophages actively eliminate transformed cells, cellular debris, pathological aggregates, and invading pathogens^2,3^. Phagocytosis begins with the engagement of phagocytic particles with specific cell surface receptors such as Fc receptors (FcR) that binds to Fc region of opsonizing IgG^1,2^. Activation of phagocytic receptors triggers actin polymerization that restructures the plasma membrane into cup-like configurations (phagocytic cups) around the particles^4,5^. Expansion of phagocytic cups leads to sequestration of particles within vesicular phagosomes. These nascent phagosomes mature into acidic, hydrolytic enzyme-rich phagolysosomes to degrade and recycle engulfed particles^6^. Components of the degraded materials are also cross-presented as antigens to activate adaptive immunity^7^. Being so critical for organismal homeostasis, disease defense and immunity, aberrant phagocytosis can be detrimental. Uncontrolled phagocytosis leads to excessive RBC destruction (hemophagocytosis), tissue damage, and neurodegeneration^8,9^; while impaired clearance of phagocytic bodies develops autoimmune, immunodeficiency and neuroinflammatory diseases^10,11^. Hence, phagocytosis requires intense control over uptake and clearance to thwart adverse consequences. Machineries that promote or inhibit phagocytosis and understanding their mechanisms are thus extremely critical^12^.

Abundance of macrophage surface receptors, expansion of phagocytic cups, and maturation of phagosomes determine the extent of particle engagement, engulfment and clearance^2,13^. Regulated fusion of FcR carrying recycling compartments to plasma membrane delivers surface receptors including FcR^2,14^. Acquisition of membranes for phagocytic cup expansion (for phagosome biogenesis)^2^ and subsequent maturation of phagosomes into degradative phagolysosomes also requires regulated fusion with various endolysosomal compartments^2,15,16^. In eukaryotes, soluble N-ethylmaleimide-sensitive factor attachment protein receptor (SNARE) proteins mediate membrane fusions^17,18^. Typically, two target membrane-localized t-SNAREs (syntaxins and SNAPs) combine with one vesicle-associated v-SNAREs (like VAMPs) to form a four-helix trans-SNARE complex (SNARE-pin) that mediates membrane merger^17,18^. SNARE-dependent trafficking of early endosomes^19^, recycling endosomes^20,21^, late endosomes^22^ and lysosomes^23,24^ promote phagocytosis. However, machineries and mechanisms that coordinate endolysosomal trafficking to balance phagocytic uptake (phagosome biogenesis) and phagolysosomal clearance remain largely unknown.

Contrary to pro-fusion roles, some SNAREs (inhibitory-/i-SNAREs) interfere with other cis-SNAREs in forming fusogenic SNARE-pin to fine-tune membrane fusions^25,26^. Syntaxin-2 (Stx2) was initially discovered as an extracellular epithelial morphogen called epimorphin^27,28^, and later its intracellular version was characterized as a t-SNARE^29^. Stx2 is unique amongst all the known t-SNAREs in showing fusion-neutral (in platelets)^30^, fusion-competent (in kidney and goblet cell)^31,32^, and i-SNARE properties (in pancreatic acinar and beta cells)^33–35^. Considering Stx2’s variegated roles in membrane fusions, coupled with prominent expression in macrophages^36–38^ but unknown functions in phagocytosis, we hypothesized that Stx2 might be a crucial component that can differentially control the delivery of diverse intracellular compartments to shape the course of phagocytosis.

We found that Stx2 is indeed a critical regulator of macrophage endomembrane trafficking, essential for controlling phagocytic uptake and phagosome maturation. Stx2 inhibits acquisition of early endosomes and VAMP4 positive post-Golgi vesicles to restrain uncontrolled expansion of phagocytic cups, hence phagosome biogenesis. Stx2 also restricts recycling of receptor carrying vesicles to curtail surface FcR expression. In contrast, Stx2 promotes phagosome maturation by the acquisition of late endosomes and lysosomes. Stx2 also facilitates macrophage secretion of pro-cathepsins to curb cellular lysosome contents. Therefore, Stx2 depleted macrophages lose control over phagocytosis and show unabated particle uptake but impaired degradation, despite having more lysosomes. Thus, our study identifies Stx2 as a novel regulator of phagocytosis and elucidates its mechanism for balancing phagocytic uptake and clearance.

## RESULTS

### Stx2 limits macrophages to engage and uptake of IgG-opsonized particles

To understand functions of Stx2 in macrophage phagocytosis, we first assessed the localization of endogenous Stx2 within the subcellular compartments of RAW 264.7 macrophages. Using confocal immunofluorescence imaging, we detected predominant presence of Stx2 on plasma membrane (Figure 1A) and on Rab5A positive early endosomes (Figure 1B; r=0.625). We also observed some moderate colocalization with Rab11-positive recycling endosomes (Figure 1C; r=0.336) and VAMP4-positive compartments (Figure 1D; r=0.411) that include trans-Golgi network, sorting and recycling endosomes^39,40^. A moderate co-localization was also observed with VAMP7-positive late endosomes (Figure 1E; r=0.304). Stx2 was localized poorly with lysosome marker protein LAMP1 (Figure 1F; r=0.154). In accordance to an earlier report^36^, we also obtained abundant presence of Stx2 on IgG-opsonized zymosan particle (OPZ) containing phagosomes (Figure 1G). Collectively, these data demonstrate Stx2’s versatile intracellular distribution in macrophages.

**Figure 1.**
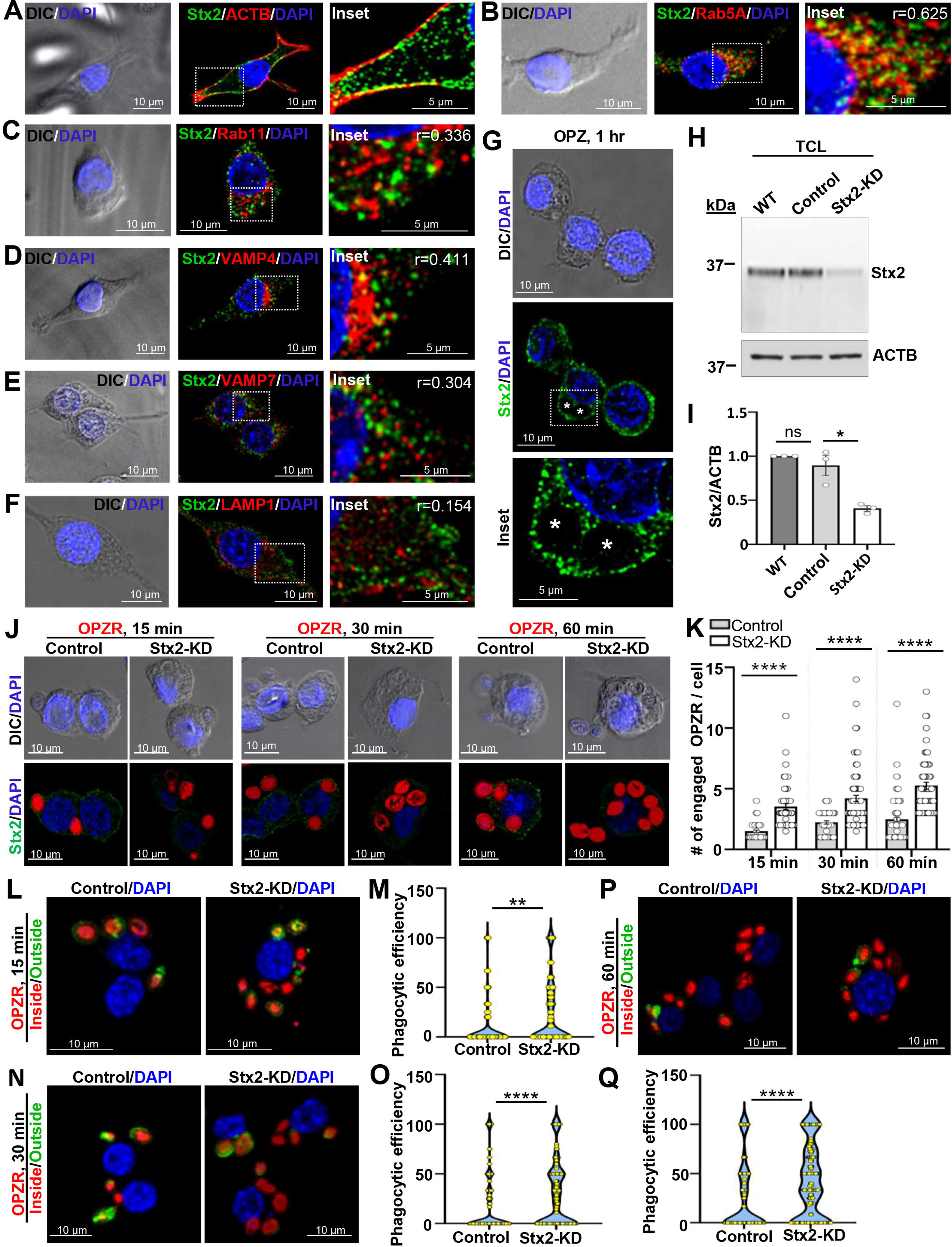
Stx2 exhibits multiple subcellular distributions in macrophages, and inhibits engagement and uptake of IgG-opsonized particles. (A-G) DIC and confocal images of endogenous Stx2 with endogenous (A) β-actin/ACTB, (B) RAB5A, (C) Rab11, (D) VAMP4, (E) VAMP7, (F) LAMP1 and with (G) OPZ containing phagosomes (white stars) in RAW 264.7 macrophages. Boxed areas enlarged in the insets. “r” values show degree of colocalization. Scale bars, 10 and 5 μm (insets). DAPI denotes the nucleus. (H) Western blot of Stx2 and β-actin (ACTB, loading control) from RAW 264.7 macrophages (WT) and macrophages transduced with scrambled (sc) shRNA (Control) or shRNAs targeted to Stx2 (Stx2-KD). (I) Quantified Stx2 band density. N = 3 independent experiments. Error bars, ± SEM. *p < 0.05, ns = not significant. (J) DIC and stacked (3 consecutive optical sections) confocal images from 15, 30 and 60 min OPZR incubated macrophages that also probed for Stx2. DAPI marks the nucleus. Scale bars, 10 μm. (K) Quantification of engaged OPZR per macrophage (15 min: Control, n= 61; Stx2-KD, n= 82; 30 min: Control, n= 56; Stx2-KD, n= 109; 60 min: Control, n= 111; Stx2-KD, n= 86). N = 3 independent experiments. Error bars, ± SEM. ****p < 0.0001. (L-Q) Maximum intensity projection confocal images of inside-outside stained macrophages incubated with OPZR for (L) 15 min, (N) 30 min or (P) 60 min, and quantification of phagocytosis efficiency for 15 min (M) (Control, n= 75; Stx2-KD, n= 68), 30 min (O) (Control, n= 273; Stx2-KD, n= 344) and 60 min (Q) (Control, n= 165; Stx2-KD, n= 190). N = 3 independent experiments. Error bars, ± SEM. **p < 0.01, ****p < 0.0001. See also Figure S1.

To evaluate Stx2 functions in macrophage phagocytosis, we established a Stx2 depleted (Stx2-KD: ∼70%) RAW 264.7 cell line by stably expressing shRNAs targeted to mouse Stx2 (Figure 1H and I; Figure S1A and B). A parallel Control cell line was generated by expressing nonspecific scrambled-shRNA (sc-shRNA) that did not affect expression of endogenous Stx2 (Figure 1H and I). In order to examine impact of Stx2 depletion on phagocytosis, we initially incubated Control and Stx2-KD macrophages with IgG-opsonized fluorescent polystyrene beads (OPFB-2) for 15 to 60 min (Figure S1C). Differential interference contrast (DIC) and confocal imaging showed increased association of OPFB-2 to Stx2-KD macrophages at all the investigated time points (Figure S1C and Figure S1D). To assess if this is a general phenomenon for IgG-opsonized particles, we exposed macrophages to opsonized zymosan red-fluorescent particles (OPZR)^23^. We also immunostained macrophages for Stx2 to directly correlate the change in particle association with Stx2 level (Figure 1J). We observed a consistent surge in OPZR engagement in Stx2 depleted macrophages (15 min: ∼2.4 fold; 30 min: ∼1.9 fold and 60 min: ∼2.2 fold) (Figure 1J and K), which are comparable to OPFB-2 (Figure S1C and D). Stx2-KD thus seemed to augment macrophage capacity for opsonized particle engagement.

To verify, whether the increased engagement translated into increased phagocytic uptake, we compared phagocytosis efficiency amongst Control and Stx2-KD RAW 264.7 macrophages exposed to OPZR for 15 to 60 min (Figure 1L-Q). An inside/outside staining approach was adopted to discriminate internalized particles from external particles^41^. Quantification revealed prominent increase in phagocytic uptake in Stx2 depleted macrophages throughout the incubation period (15 min: ∼2.9 fold; 30 min: ∼1.69 fold and 60 min: ∼1.65 fold) (Figure 1L-Q). Thus, in addition to increased particle engagement, Stx2 depletion also augments macrophage capacity for phagocytic uptake.

### Enhanced formation and expansion of phagocytic cups in Stx2 depleted macrophages

Engagement of IgG-opsonized particles with macrophages develops phagocytic cups for grasping phagocytic particles^2,5^. Expansion of phagocytic cups sequesters particles within vesicular phagosomes^2,5^. Stx2-KD enhances the engagement and uptake of IgG-opsonized particles (Figure 1J-Q). We therefore utilized scanning electron microscopy (SEM) to scrutinize alterations in macrophage surface morphology during phagocytosis of IgG-opsonized polystyrene beads of ∼3.0 µm diameter (OPB-3). At an earlier time point (5 min) Stx2-KD macrophages exhibited ∼2 fold increment in OPB-3 attachment (Figure 2A and C), with clear emergence of phagocytic cups (Figure 2A and B). The numbers of attached OPB-3 were further increased by ∼1.8 fold after 10 min, accompanied by prominent appearance of phagocytic cups (Figure 2A and B). By 30 min, there was a marked decrease in attached OPB-3 in both Control and Stx2-KD macrophages, with the majority of the attached OPB-3 internalized (Figure 2A and C). For a quantitative assessment of phagocytic cup expansion, we measured the individual OPB-3 area (in µm^2^) that covered with plasma membrane (Figure 2D and E). Analysis demonstrated ∼1.8 fold increment over 5 min and ∼2.67 fold increment over 10 min and ∼2.33 fold increase over 30min (Figure 2D). Analysis of cumulative bead area covered in individual macrophage showed ∼2.17 fold increment over 5 min, ∼3.56 fold increment over 10 min and ∼3.87 fold increase at 30 min in Stx2-KD macrophages (Figure 2E). Collectively, these data suggest that Stx2 depleted macrophages can expand plasmalemma uncontrollably for particle trapping and uptake.

**Figure 2.**
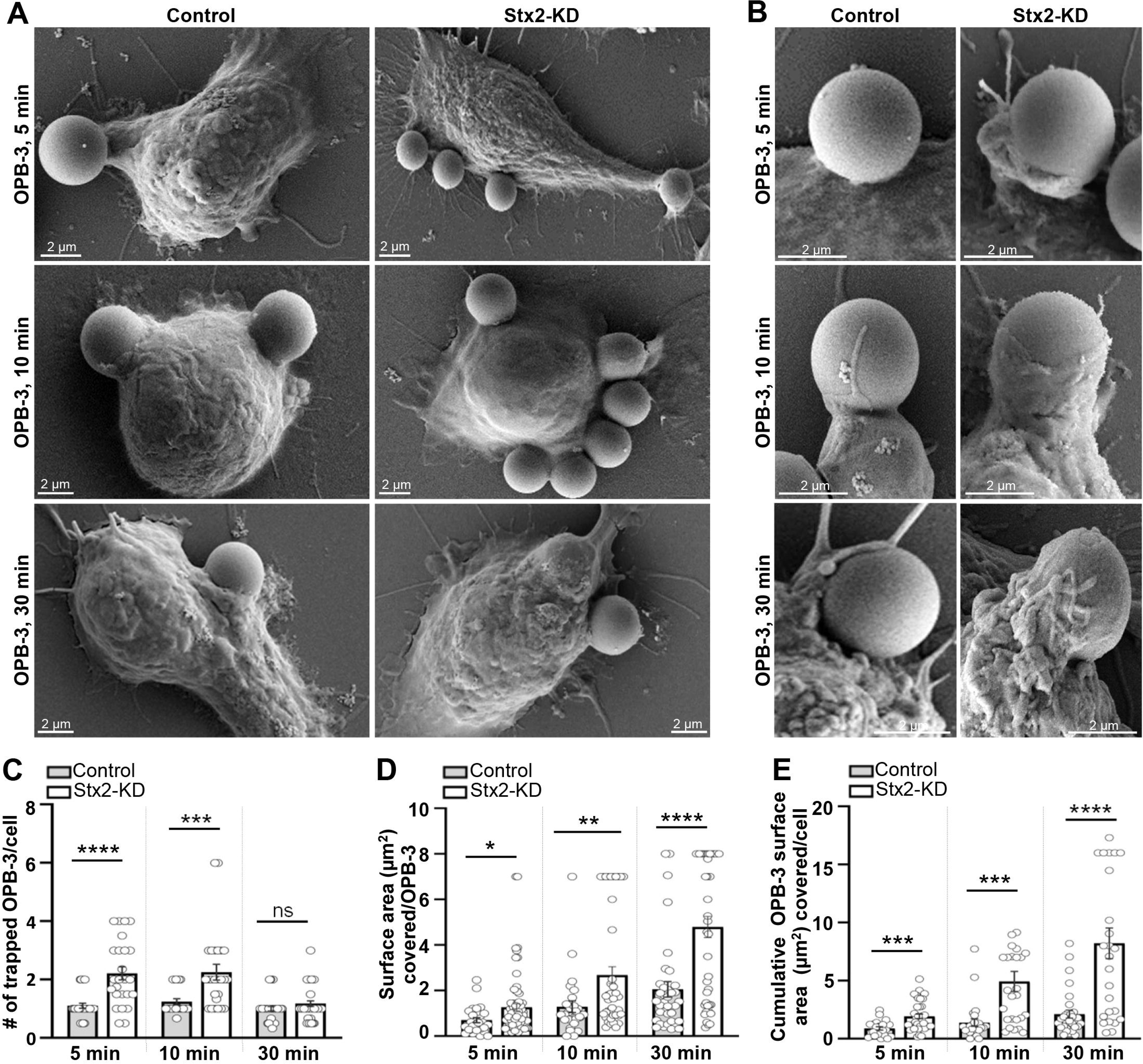
Enhanced formation and expansion of phagocytic cups in Stx2 depleted macrophages. (A) Representative scanning electron micrographs of RAW 264.7 macrophages incubated with OPB-3 for 5, 10 and 30 min. Scale bars, 2 μm. Additional set of images in Figure S2. (B) Magnified micrgraphs show single phagocytic cup. Scale bars, 2 μm. (C) Quantification of the surface attached OPB-3 per macrophage (5 min: Control, n= 29; Stx2-KD, n= 25; 10 min: Control, n= 23; Stx2-KD, n= 25; 30 min: Control, n= 33; Stx2-KD, n= 36). N = 3 independent experiments. Error bars, ± SEM. ns = not significant, ***P < 0.001, ****P < 0.0001. (D and E) Quantification of the (D) mean surface area (µm^2^) covered in individual OPB-3 and (E) cumulative bead surface areas (µm^2^) wrapped with plasma membrane (5 min: Control, n= 17; Stx2-KD, n= 22; 10 min: Control, n= 20; Stx2-KD, n= 17; 30 min: Control, n= 19; Stx2-KD, n= 23). N = 3 independent experiments. Error bars, ± SEM. ns = not significant, *P < 0.05, **P < 0.01, ***P < 0.001, ****P < 0.0001.

### Stx2 curbs surface-expressed FcR by controlling receptor recycling

Phagocytic receptors engage phagocytic particles with phagocytes^4^. Stx2 depleted macrophages show heightened engagement of IgG-opsonized particles (Figure 1J). We therefore assessed surface expression of IgG Fc region binding phagocytosis receptors FcR. Non-permeabilized macrophages were immunostained with FcR-specific antibodies to label surface receptors and subjected to FACS analysis that showed ∼2.3 fold increment in Stx2-KD macrophages (Figure 3A and B). Consistently, non-permeabilized Stx2-KD macrophages produced ∼1.7 fold higher fluorescence intensity for FcR in confocal imaging (Figure 3C and D). To address whether this increased surface FcR abundance is an outcome of increased cellular expression or increased plasmalemmal delivery, we assessed total FcR content in permeabilized macrophages by FACS that showed rather slight drop in Stx2 depleted macrophages (Figure 3E and F). Surface delivery of FcR relies on recycling from its intracellular endosomal stores^42^. We therefore challenged the macrophages with OPZ for 15 min and FACS-analyzed the surface FcR content (Figure 3G). Interestingly, despite higher phagocytic efficiency (hence more internalization of FcR), the surface level of FcR remains comparable to unchallenged Stx2-KD macrophages and Control macrophages (Figure 3G). We also obtained higher surface expression of transferrin receptor (TfR) despite having no change in total cellular expression (Figure 3H-K). FcR and TfR take same recycling route to plasma membrane, controlled by Rab11^14^ that also show higher expression in Stx2-KD macrophages (Figure 3L-O). Thus, suggests increased recycling of FcR for increased surface FcR for increased particle engagement.

**Figure 3:**
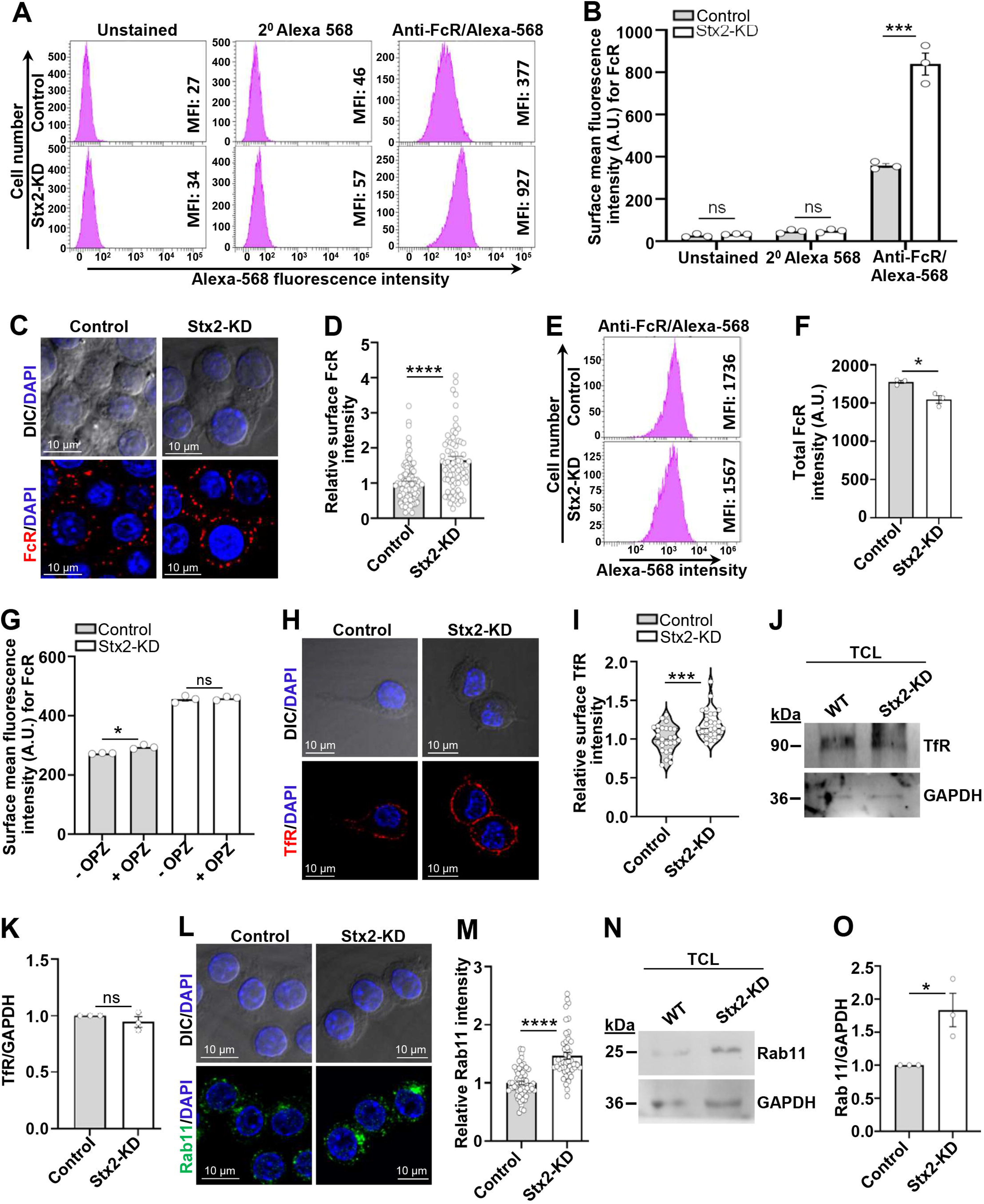
Stx2 curbs surface-expressed FcR by controlling receptor recycling. (A) FACS histograms with mean fluorescence intensity (MFI) of unlabeled, only 2^0^-antibody labeled and FcR labeled macrophages. (B) Mean MFI values of 3 independent FACS histograms. Error bars, ± SEM. ns = not significant, ***P < 0.0001. (C) DIC and confocal images of surface FcR. DAPI marks the nucleus. Scale bars, 10 μm. (D) Quantified surface fluorescence intensity for FcR (Control, n= 105; Stx2-KD, n= 78). N = 3 independent experiments. Error bars, ± SEM. ****P < 0.0001. (E) FACS histograms for total FcR with MFI values. (F) Mean MFI values for total FcR from 3 independent FACS histograms. Error bars, ± SEM. ns = not significant, *P < 0.05. (G) Mean MFI values of surface FcR from 3 independent FACS histograms of OPZ fed (15 min) macrophages. Error bars, ± SEM. ns = not significant, *P < 0.05. (H) DIC and confocal images of surface TfR. Nuclei were labeled with DAPI. Scale bars, 10 μm. (I) Quantified surface TfR fluorescence intensity (Control, n= 27; Stx2-KD, n= 27). N = 3 independent experiments. Error bars, ± SEM. ***P < 0.001. (J) Western blot of TfR and GAPDH (loading control) from macrophages. (K) Quantified TfR band density normalized to GAPDH. N = 3 independent experiments. Error bars, ± SEM. ns = not significant. (L) DIC and confocal images for Rab11. DAPI shows the nuclei. Scale bars, 10 μm. (M) Quantified Rab11 fluorescence intensity (Control, n= 59; Stx2-KD, n= 49). N = 3 independent experiments. Error bars, ± SEM. ****P < 0.0001. (N) Western blots of Rab11 and GAPDH (loading control) from macrophages. (O) Quantified Rab11 band density. N = 3 independent experiments. Error bars, ± SEM. *P < 0.05.

### Stx2 prevents non-lytic compartment acquisition for phagocytic cup expansion and phagosome biogenesis

Expansion of phagocytic cups for phagosome biogenesis requires adequate supply of membranes from intracellular reservoirs^2,39^. We next aimed to identify and compare the contributions of specific intracellular compartments contributing in phagosome biogenesis during early point of uptake (Figure 4A-F). Purified nascent phagosome enriched fractions from briefly (30 min) OPB-0.8 exposed macrophages revealed ∼1.7 fold increased concentration of early endosome protein Rab5A (Figure 4A and B) and ∼1.43 fold increased content of VAMP4 in Stx2-KD macrophage derived phagosomes (Figure 4A-C). VAMP4 has been shown to traffic through recycling and sorting endosomes to enrich trans-Golgi network, which is known to be absent from FcR-induced phagosomes^43,44^. Consistently, we could not detect trans-Golgi dwelling golgin-160 (Figure 4A and D), suggesting contributions from VAMP4-positive post-Golgi compartments. Interestingly, contributions from late endosomes (VAMP7)^22^ and lysosomes (VAMP8)^45^ were depleted by 1.42 fold and 2.9 fold (Figure. 4A, E and F) respectively.

**Figure 4:**
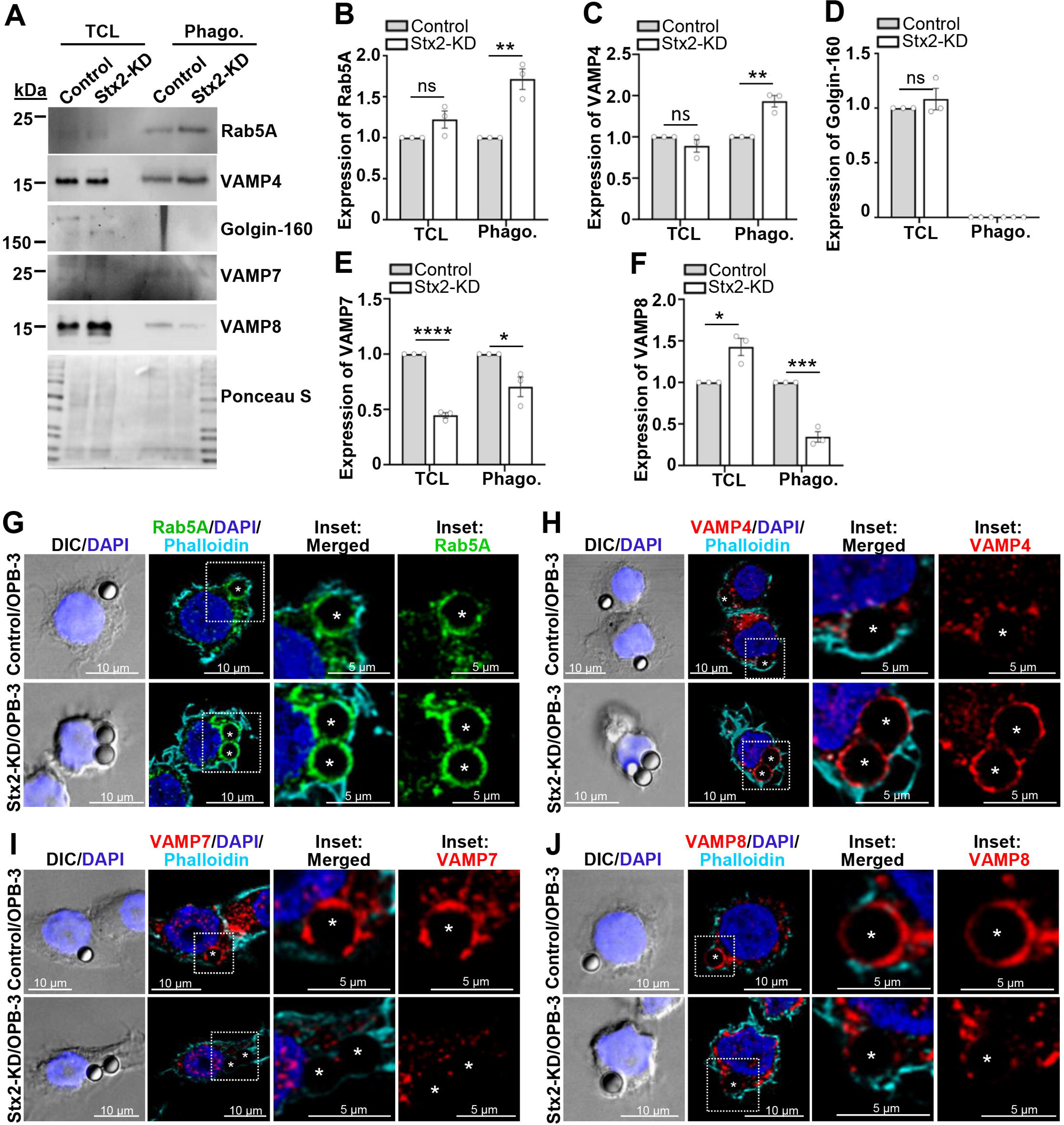
Stx2 prevents delivery of early endosomes and VAMP4-positive post-Golgi compartments for phagocytic cup expansion/phagosome biogenesis. (A) Western blots of macrophage lysates (TCL) and 30 min OPB-0.8 fed purified phagosomes (phago), probed for Rab5A, VAMP4, Golgin-160, VAMP7 and VAMP8. Ponceau S stained membrane shows uniform loading. (B-F) Quantified band density of (B) RAB5A, (C) VAMP4, (D) Golgin-160, (E) VAMP7 and (I) VAMP8. N = 3 independent experiments. Error bars, ± SEM. ns = not significant, *P < 0.05, **P < 0.01, ***P < 0.001, ****P < 0.0001. (G-J) DIC and confocal images of 30 min OPB-3 challenged macrophages for F-actin (phalloidin) and (G) Rab5A, (H) VAMP4, (I) VAMP7 and (J) VAMP8. DAPI shows the nuclei. White stars mark OPB-3 containing phagosomes. Boxed regions enlarged in the insets. Scale bars 10 and 5 μm (inset).

To verify and visualize the acquisition of these compartments, we imaged OPB-3 engulfing macrophages (Figure 4G-J). Consistent to the purified phagosomes (Figure 4A-F), both Rab5A (Figure 4G) and VAMP4 (Figure 4H) showed increased coalescence around the nascent phagosomes (containing F-actin) of Stx2-KD macrophages, while the staining appeared punctate and incomplete (phagocytic cups) for Control macrophages. Similar staining with VAMP7 (Figure 4I) and VAMP8 (Figure 4J) showed their reduced presence around the nascent phagosomes and phagocytic cups of Stx2-KD macrophages. Altogether, Stx2 curtails delivery of non-lytic early endosomes and VAMP4-positive post-Golgi compartments for phagocytic cup expansion and phagosome biogenesis.

### Increased lysosome content and diminished pro-cathepsin secretion in Stx2 depleted macrophages

Intrigued by the heightened expression of VAMP8 (Figure 4A and F), a lysosomal SNARE protein^45^, we postulated that reduction of Stx2 might also augment lysosome content in macrophages. To investigate, we initially stained macrophages with LysoTracker Red DND-99 (that labels acidic endolysosomes), and subjected them to FACS analysis (Figure 5A). We observed ∼1.6 fold increase in the MFI in Stx2 reduced macrophages (Figure 5A). Additionally, confocal imaging disclosed notable increase in both abundance and intensity (∼1.8 fold higher MFI) of red fluorescent puncta on Stx2 depletion (Figure S3A and B). These findings indicate sprawling presence of acidic endolysosomes in Stx2 depleted macrophages. We then immunostained macrophages for lysosome membrane protein LAMP1 to directly visualize lysosomes that rose by ∼1.7 fold (Figure 5B and C). Further, western blot analysis validated ∼2 fold Increased abundance of LAMP1 in Stx2 depleted macrophages (Figure S3C and D). Expression of major lysosomal hydrolases^46^ cathepsin L (CTSL), cathepsin B (CTSB), and cathepsin D (CTSD) were also higher in Stx2 depleted macrophages (Figure 5D-G). Western blots confirmed increase in both premature (pro-cathepsins) and catalytically active mature cathepsins (Figure 5F and G). An elevated basal autophagic flux, as evidenced by heightened LC3-II/LC3-I ratio and degradation of autophagy substrate P62/SQSTM1 (Figure S3E-G) further attests increase in functional and fusion competent lysosomes. In order to explain increased lysosome contents, we examined the status of transcription factor EB (TFEB), the master regulator for lysosome biogenesis^47,48^. TFEB expression was similar in both Control and Stx2-KD macrophages (Figure 5H and S3H). However, TFEB was undetectable in nuclear fraction and remained confined in the cytosol (Figure 5H and S3H), suggesting no involvement. Interestingly, secretion assay for cathepsins revealed specific reduction of pro-cathepsins in the macrophage culture media (Figure 5I-K). We hypothesize that increased retention of pro-cathepsins can lead to increased transition to mature cathepsin containing lysosomes to accumulate within Stx2 depleted macrophages.

**Figure 5:**
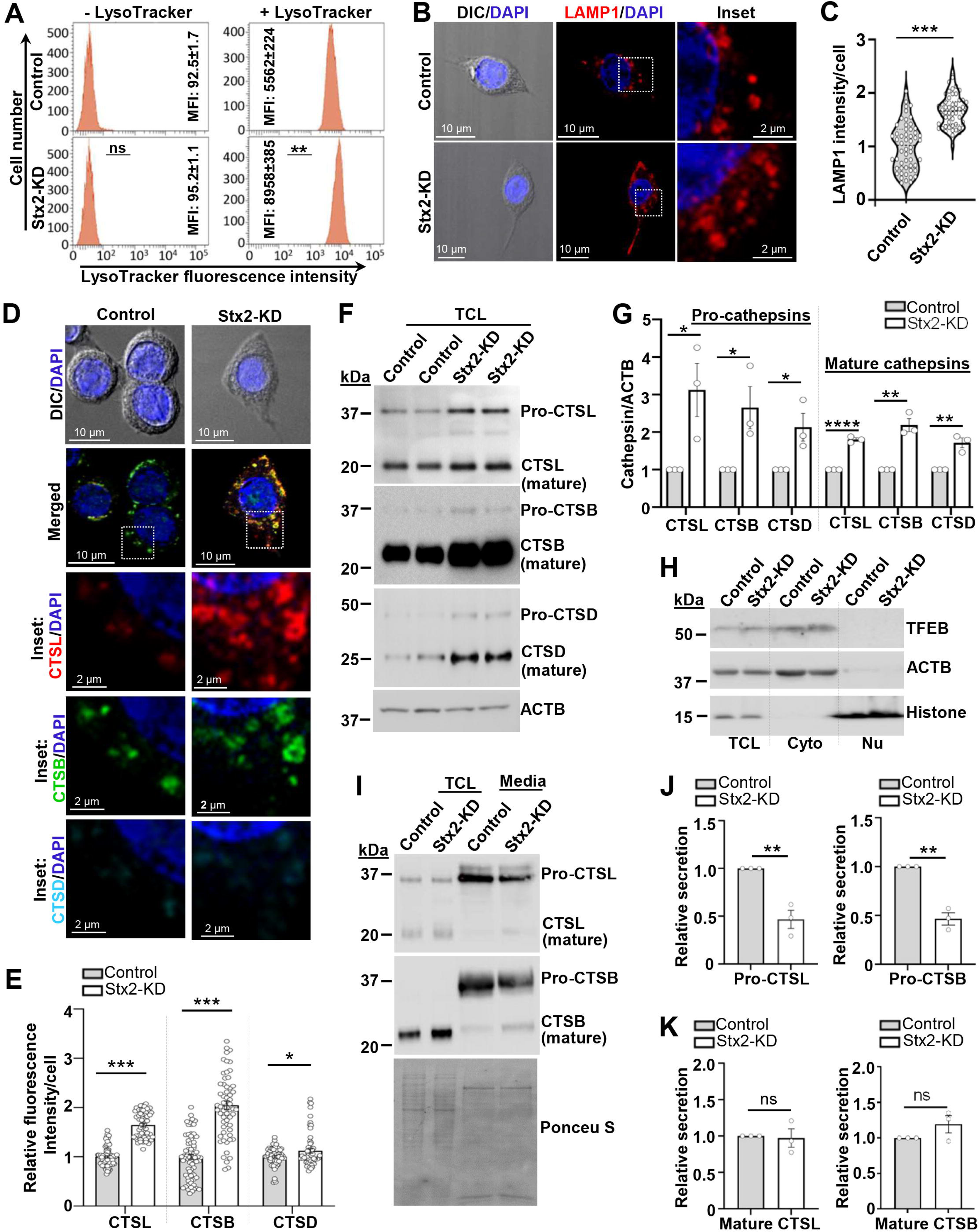
Increased lysosome content couples with diminished pro-cathepsin secretion in Stx2 depleted macrophages. (A) FACS histograms and the mean MFI values (as ± SEM) of 3 histograms from LysoTracker stained RAW 264.7 macrophages. ns = not significant, **P < 0.01. (B) DIC and confocal images of LAMP1 immunostained macrophages. Boxed areas enlarged in inset. Nuclei labeled with DAPI. Scale bars 10 and 2 µm (inset). (C) Quantified LAMP1 fluorescence intensity (Control, n= 54; Stx2-KD, n= 54). N = 3 independent experiments. Error bars, ± SEM. ***P < 0.001. (D) DIC and confocal merged images for CTSL, CTSB and CTSD. Boxes are magnified and split into separate channels in inset. Scale bars 10 and 2 μm (inset). (E) Quantified (Control, n= 73; Stx2-KD, n= 56) fluorescence intensity for cathepsins. N = 3 independent experiments. Error bars, ± SEM. *P < 0.05, ***P < 0.001. (F) Western blots of macrophages for CTSL, CTSB, CTSD and ACTB (loading control). (G) Quantified band density for CTSL, CTSB and CTSD. N = 3 independent experiments. Error bars, ± SEM. **P < 0.01, ***P < 0.001. (H) Western blot detection for TFEB in nuclear and cytosolic fractions. N=3 independent experiments. (I) Western blot detection of macrophage secreted pro- and mature-CTSL and CTSB. Ponceau S stained membrane shows equal loading. (J-K) Quantified secreted (J) pro-CTSL (left) and pro-CTSB (right) and (K) mature-CTSL (left) and mature-CTSB (right). N = 3 independent experiments. Error bars, ± SEM. ns = not significant. **P < 0.01. See also Figure S3.

### Stx2 is required for phagosome maturation by acquisition of late endosomes/lysosomes and phagosome acidification

Extensive abundance of phagosomes (Figure 1) and lysosomes (Figure 5) but reduced delivery of late endosomes and lysosomes to budding phagosomes in Stx2 depleted macrophages (Figure 4I and J) prompted us to investigate possible defect in phagosome maturation. To delineate, we exposed macrophages to OPZ for 15 to 60 min (Figure 6A). In line with our prior observations (Figure 1), Stx2-KD macrophages engulfed more OPZ particles throughout the experimental timeframe (Figure 6A). We observed a gradual and prominent increase of LAMP1 fluorescence intensity around the Control phagosomes, compared to Stx2 depleted macrophages (Figure 6A and B). Concurrently, the presence of CTSL, CTSB and CTSD also remained low in Stx2 depleted phagosomes (Figure 6C and D). These observations were validated using purified phagosomes from longer OPB-0.8 incubated (1 h uptake and 30 min maturation) macrophages. We obtained reduced amount of LAMP1 as well as catalytically active mature CTSL, CTSB and CTSD in Stx2 depleted phagosomes (Figure 6E and F). Interestingly, despite comparable presence of premature CTSD (Figure 6E), the mature CTSD was very low in Stx2 depleted phagosomes (Figure 6E and I). Since CTSD maturation depends on CTSL and CTSB activity^49^, our observations further suggest that in addition to their reduced phagosomal presence, the proteolytic activity of CTSL and CTSB were also low in Stx2-KD phagosomes (Figure 6E and G-H). As cathepsins remain active in acidic pH^50^, we additionally probed for phagosome acidifying vacuolar ATPase^51^ subunit ATP6V1A, which showed reduced presence in Stx2 depleted phagosomes (Figure 6E and J), suggesting reduced phagosome acidification. To confirm this we allowed macrophages to engulf opsonized pHrodo red zymosans (OpHRZ), whose fluorescence increases with reduction in pH (Figure 6K). Internalization of OpHRZ was confirmed by inside (red)/outside (green) staining. Internalized OpHRZ particles in Control cells produced bright red fluorescence upon 30 min that further increased at 60 min (Figure 6K and L). However, OpHRZ particle fluorescence intensity remained consistently dim within Stx2-KD macrophages (Figure 6K and L). The MFI of the OpHRZ particles were also remained significantly low in both 30 min and 60 min (Figure 6L). Based on these findings, we conclude that Stx2 is an essential for maturation and acidification of phagosomes in macrophages.

**Figure 6.**
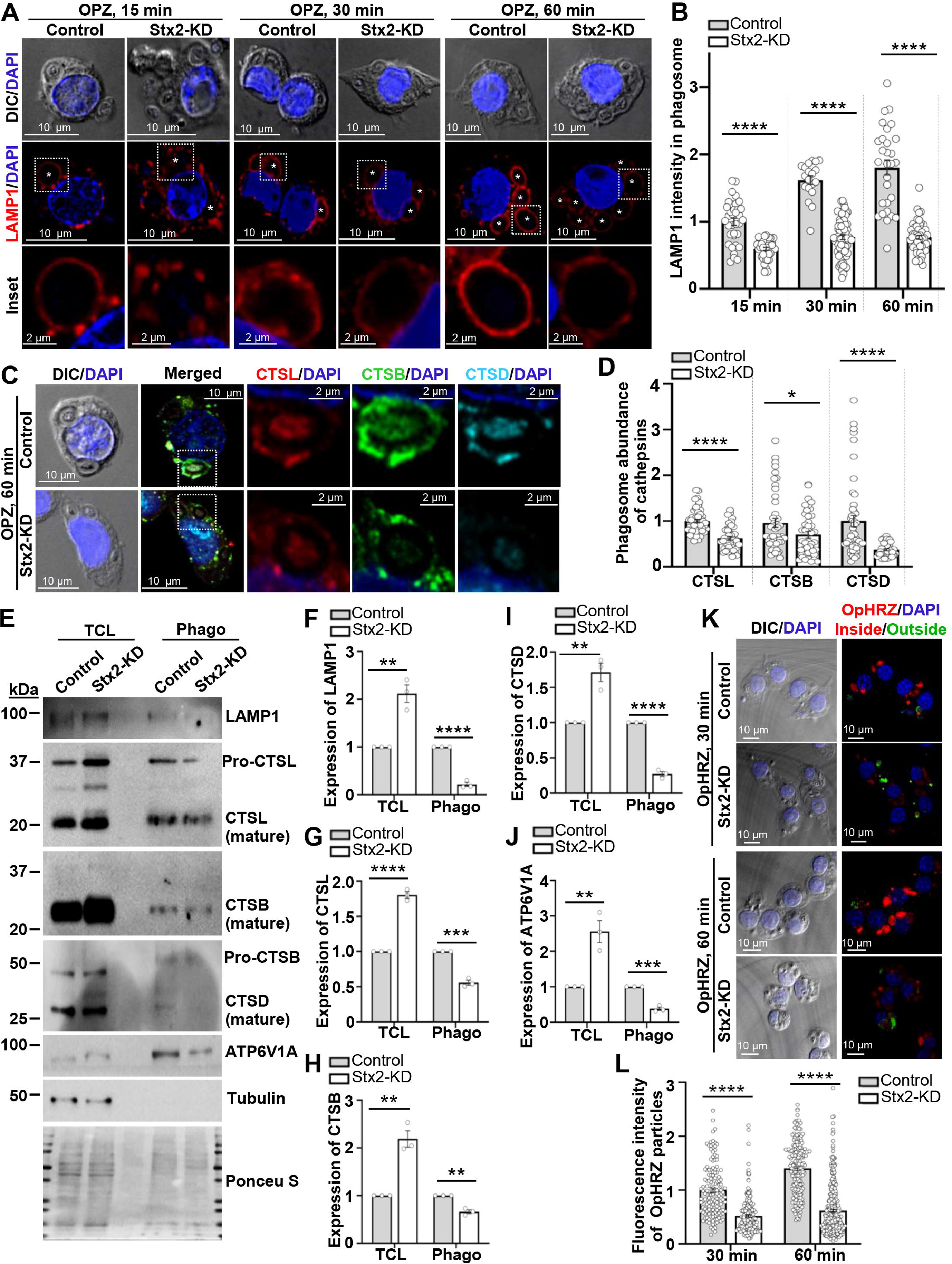
Stx2 is required for macrophage phagosome maturation and acidification. (A) DIC and confocal images of OPZ fed macrophages immunostained for LAMP1. White stars show the OPZ-containing phagosomes. Boxed regions magnified in the inset. DAPI marks the nuclei. Scale bars 10 and 2 μm (inset). (B) Quantification (15 min: Control, n= 35; Stx2-KD, n= 42; 30 min: Control, n= 29; Stx2-KD, n= 68; 60 min: Control, n= 29; Stx2-KD, n= 53) of the phagosomal LAMP1 fluorescence intensity. N = 3 independent experiments. Error bars, ± SEM. ns = not significant. ****P < 0.0001. (C) DIC and confocal images of 60 min OPZ fed macrophages immunostained for CTSL, CTSB, and CTSD. Boxed areas are magnified and shown as separate channels in inset. DAPI labeled the nuclei. Scale bars 10 and 2 μm (inset). (D) Quantification (Control, n= 46; Stx2-KD, n= 43) of the phagosomal fluorescence intensity for CTSL, CTSB and CTSD. N = 3 independent experiments. Error bars, ± SEM. *P < 0.05, ****P < 0.0001. (E) Western blots of macrophages and purified phagosomes (phago) probed for LAMP1, CTSL, CTSB, CTSD, ATP6V1A and tubulin (loading control for macrophages and negative control for “phago”). Ponceau S stained membrane shows uniform loading. (F-J) Quantified band density of (F) LAMP1, (G) CTSL, (H) CTSB, (I) CTSD and (J) ATP6V1A. N = 3 independent experiments. Error bars, ± SEM. **P < 0.01, ***P < 0.001 and ****P < 0.0001. (K) DIC and confocal images of 30 and 60 min OpHRZ fed macrophages. DAPI used to visualize nuclei. Scale bars 10 µm. (L). Quantification of fluorescence intensity (higher intensity indicates lower pH) of internalized (red) OpHRZ particle (30 min: Control, n= 113; Stx2-KD, n= 66; 60 min: Control, n= 129; Stx2-KD, n= 131). N = 3 independent experiments. Error bars, ± SEM. ****P < 0.0001. See also Figure S4.

### Unabated uptake but impaired clearance of bacteria by Stx2 depleted macrophages

The antibacterial efficacy of macrophages relies on their ability to engulf bacteria into phagosomes and subsequently their enzymatic degradation within phagolysosomes^52^. Increased phagocytic uptake but delayed phagosome maturation on Stx2 depletion (Figure 1 and 6) led us to assess Stx2 effect on bacterial phagocytosis. We used IgG-opsonized mCherry expressing *Escherichia coli* (OPEC). Stx2 depleted macrophages trapped high numbers (∼1.86 fold) of OPEC over Control RAW cells (Figure 7A and B). FACS analysis unveiled 87.1±0.99% of Stx2-KD macrophages were associated with OPEC compared to only 45.03±2.63% Control macrophages (Figure 7C). The MFI, serving as an indicator of the degree of association of fluorescent particles per individual cell, was ∼1.91 fold higher in Stx2-KD cells (Figure 7D). Fluorescence imaging of OPEC incubated macrophages subjected to inside/outside staining revealed ∼1.38 fold rise in phagocytic uptake efficiency in Stx2-KD macrophages (Figure 7E and F). Longer incubation of OPEC (1 h internalization plus 1 h phagosome maturation) to examine macrophage digestion and clearance efficacy showed majority of the internalized OPEC became fragmented in Control macrophages (Figure 7G) while remained intact in Stx2 depleted macrophages (Figure 7G). Stx2 is thus critical in controlling bacterial clearance by phagocytosis in macrophages.

**Figure 7.**
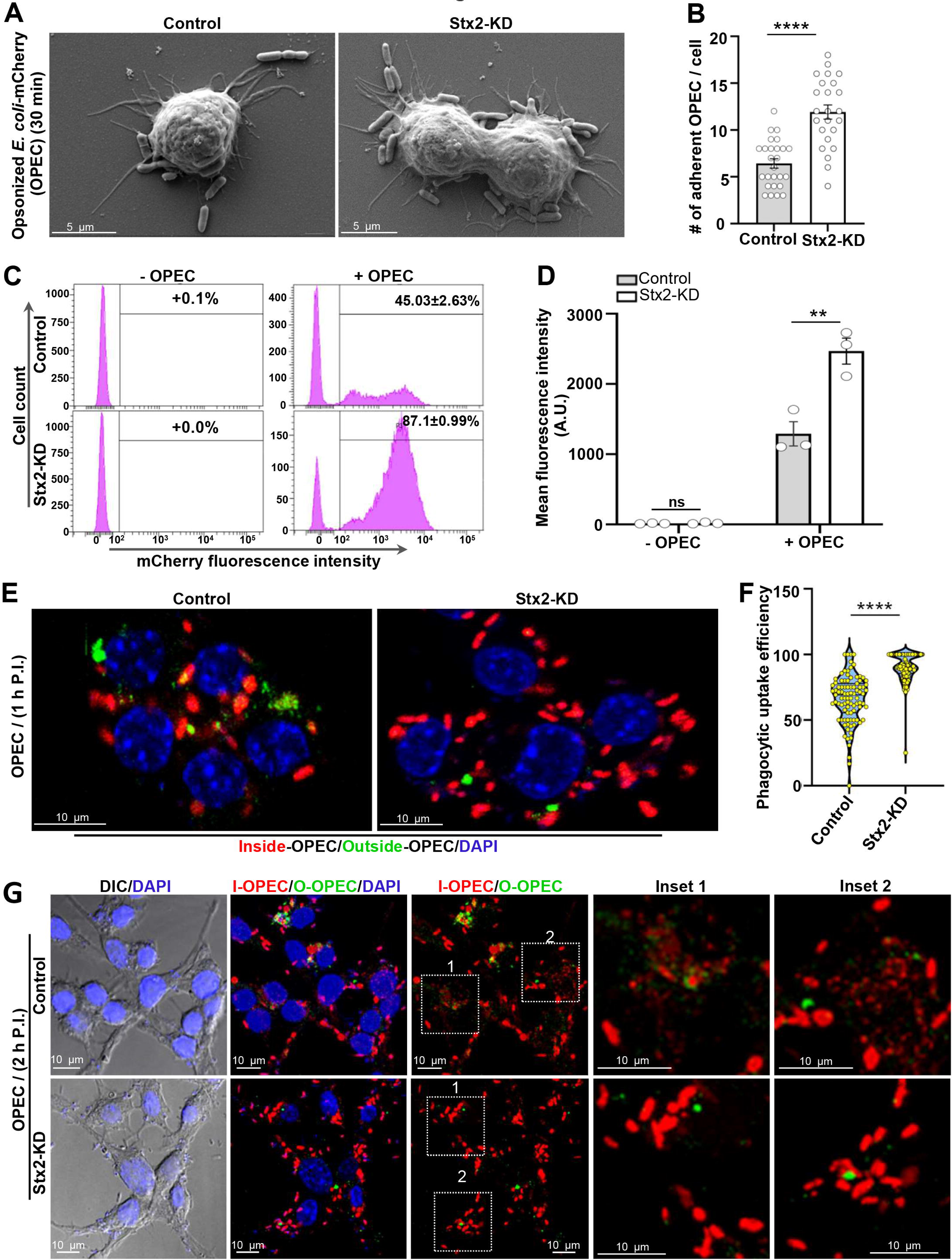
Uncontrolled uptake of IgG-opsonized *E. coli* and impaired clearance increase bacterial load in Stx2 depleted macrophages. (A) Representative SEM images of macrophages challenged with IgG-opsonized mCherry-expressing *E. coli* (OPEC) for 30 min. (B) Quantification of the surface adherent OPEC per macrophage (Control, n= 26 ; Stx2-KD, n= 24). N = 3 independent experiments. Error bars, ± SEM. ****P < 0.0001. (C) Representative FACS histograms showing extent of OPEC engagement to macrophages shown as shift in fluorescence intensity. (D) Mean MFI (OPEC/cell) from 3 independent FACS histograms. Error bars, ± SEM. Ns = not significant, **P < 0.01. (E) Representative confocal maximum intensity projection images of macrophages inside (red)/outside (green) stained for OPEC. DAPI stains the nuclei visible. Scale bars 10 µm. (F) Quantification (Control, n= 141; Stx2-KD, n= 141) of macrophage phagocytosis efficiency for OPEC. N = 3 independent experiments. Error bars, ± SEM. ****P < 0.0001. (G) Representative DIC and confocal maximum intensity fluorescence images of 2 h (1 h uptake + 1 h maturation) OPEC for incubated macrophages inside/outside stained for OPEC. Nuclei were stained with DAPI. Boxed areas are magnified in the inset. Scale bars, 10 μm, 5 µm (inset).

## DISCUSSION

In this study, we characterized the role of the SNARE protein Stx2 in macrophage phagocytosis for the very first time. We found Stx2 coordinates the trafficking of multiple intracellular compartments to balance phagocytic uptake and clearance of IgG-opsonized particles. This is corroborated with vast distribution of Stx2 on plasma membrane, early/late endosomes, VAMP4-positive compartments and phagosomes, which are directly involved in phagocytosis^2,39^. On depletion of Stx2 (Stx2-KD), we noticed aberrant engagement and uptake of IgG-opsonized particles. This enhanced engagement is attributed to higher surface expression of FcR, which we believe is an outcome of general increase in receptor recycling, as total macrophage receptors (FcR and TfR) remained similar upon Stx2-KD, but increased the content of receptor recycling GTPase Rab11^14^. Increased FcR engagement explains increased formation of phagocytic cups that requires FcR-actin axis for plasma membrane remodelling^53^.

Expansions of phagocytic cups in forming phagosomes and maturation of phagosomes into phagolysosomes require regulated acquisition of early endosomes, recycling endosomes, late endosomes and lysosomes mediated by SNAREs^2,19,20,22,23^. Stx2 depleted macrophages can uncontrollably expand phagocytic cups that accompanied with enhanced delivery of early endosomes and VAMP4-positive compartments. Previous proteomic and western blot analysis confirmed abundant presence of VAMP4 in phagosomes^39^, although its primary residence trans-Golgi networks remain abstained from IgG-opsonized particle phagocytosis^43,44,54^. Our study could not detect trans-Golgi marker protein in phagosomes either. Since VAMP4 traffics to trans-Golgi through sorting and recycling endosomes^40^, phagosomal delivery might be enhanced for these compartments. As we could not confirm this with specific markers, we kept this as “VAMP4-positive post Golgi” compartments.

Another intriguing finding from our study is that Stx2 depletion also increases macrophage lysosome contents. TFEB is the most prominent element that increases lysosomal protein synthesis for lysosome biogenesis^48^. We could not detect TFEB in nucleus, suggesting TFEB-independent mechanisms. Pro-cathepsins from trans-Golgi enroute to early endosomes that convert to late endosomes and fuse with lysosomes for full maturation of cathepsins^55^. Considering Stx2’s requirement for phagosomal hydrolase acquisition, we argued that Stx2 might also require for hydrolase secretion. A specific reduction of pro-cathepsins but not mature forms can lead to increased pro-cathepsin flux for increased lysosome biogenesis^55^. Further, an increased basal autophagy flux attests presence of catalytically active and fusion competent lysosomes. Thus, Stx2 is also critical in regulating macrophage lysosome content for catabolic homeostasis^56^.

But how Stx2 depletion can selectively upregulates acquisition of some compartments but inhibits others? Previous works on Stx2 disclosed its dual role in membrane fusions. For instance, in mucin secreting goblet cells^32^ and in rat kidney cells^31^, Stx2 pairs up with VAMP8 to mediate membrane fusions. Conversely, in zymogen-secreting pancreatic acinar cells^33^ and in insulin-secreting pancreatic beta cells^34,35^, Stx2 acts as an inhibitory (i)-SNARE by binding with VAMP8 and VAMP2, and thereby depriving other cis-syntaxins in forming fusogenic SNAREpins. Our data suggest that Stx2 acts as i-SNARE during acquiring non-lytic compartments for phagosome biogenesis and receptor recycling, but acts as fusogenic SNARE for acquiring late endosomes and lysosomes. However, the identification of partner SNAREs and the mechanisms that regulate Stx2’s contradictory behaviour need further investigation.

Nonetheless, our work unequivocally demonstrates critical involvement of Stx2 in macrophage phagocytosis. The opposing role of Stx2 in preventing excess uptake and concurrent promotion in lysosome directed clearance of engulfed particles is unique amongst all known macrophage SNAREs. This balancing act of Stx2 could have far reaching physiologic importance. Stx2 could be targeted to modulate phagocytosis plasticity for controlling hyperactive and timid phagocytosis related pathogenesis^8–11^. It would be interesting to see the status and functions of macrophage Stx2 in such pathogenesis.

## ACKNOWLEDGEMENTS

We thank DBT-Builder funded (BT/INF/22/SP45383/2022) IISER Kolkata core imaging facility. We also thank Mr. Tamal Ghosh and Mr. Kashinath Sahu for their help with FACS and for SEM, respectively. R. D. lab members are acknowledged for their technical support. We thank Dr. Arnab Gupta for providing critical reagents. We acknowledge Prof. Jayasri Das Sarma and Dr. Sankar Maiti for valuable research inputs. This work was supported by the research grants from SERB-SURE (SUR/2022/001269) and DBT Ramalingaswami Re-entry Fellowship (BT/RLF/Re-entry/07/2019) awarded to S. D., and ICMR (6/9-7(318)/2023-ECD-II) awarded to R. D.

## AUTHOR CONTRIBUTIONS

Conceptualization, S.D.; Methodology, S.S., S.D., A.N., and R.D.; Investigation, S.S., S.D., and A.N.; Formal Analysis, S.S. and S.D.; Writing – Original Draft, S.D. and R.D.; Writing – Review & Editing, All authors; Supervision and Funding Acquisition, S.D. and R.D.

## STAR*METHODS

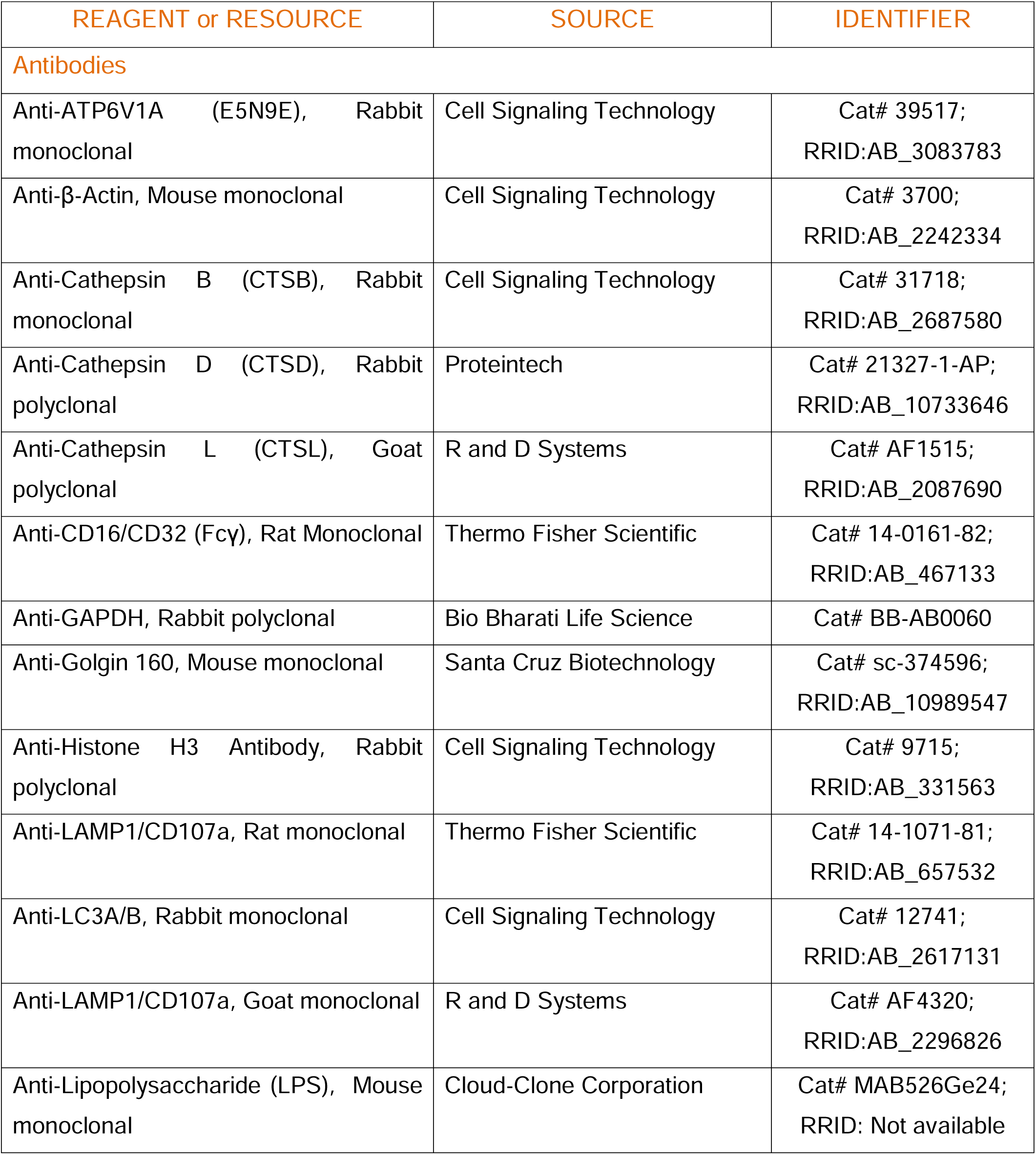
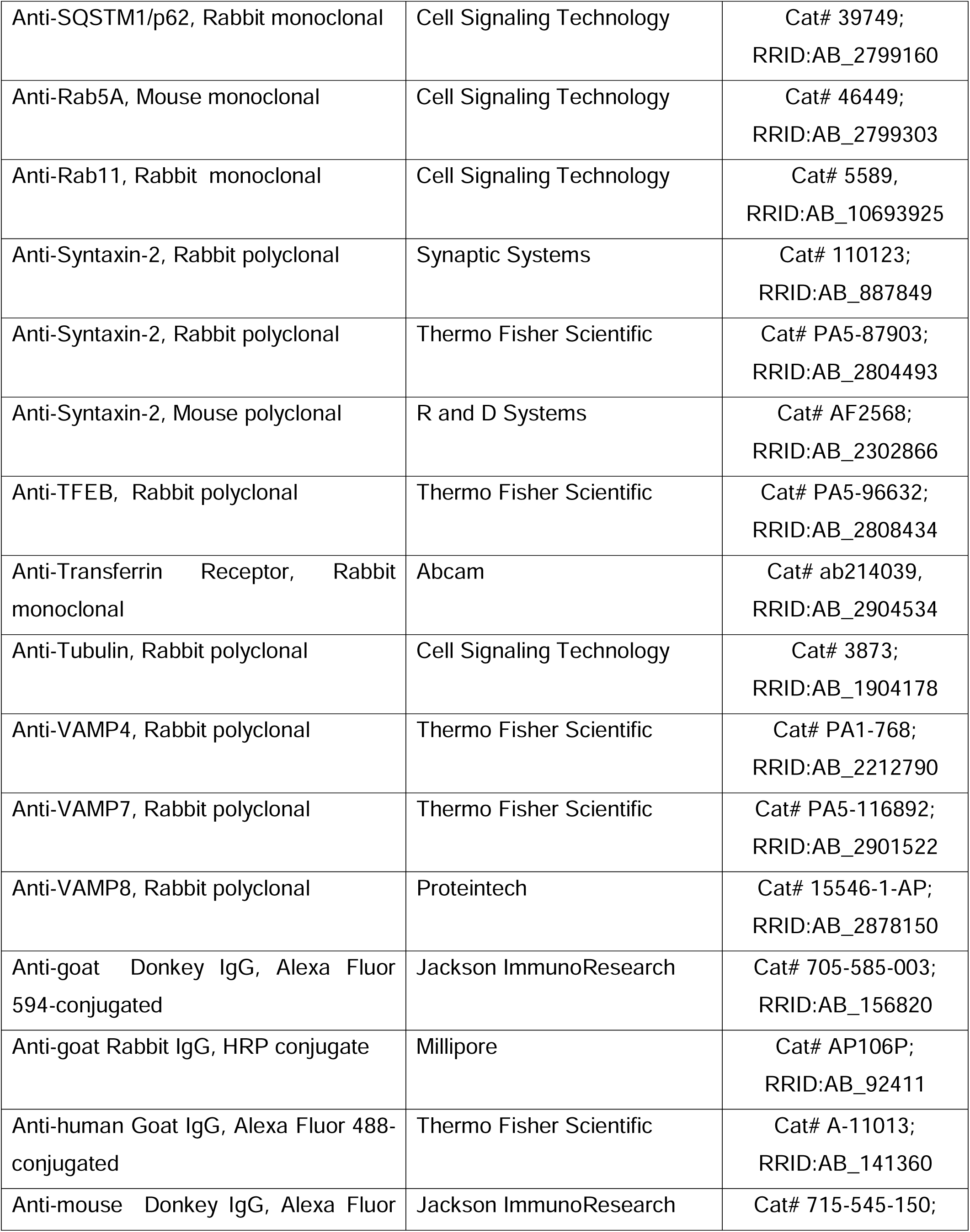
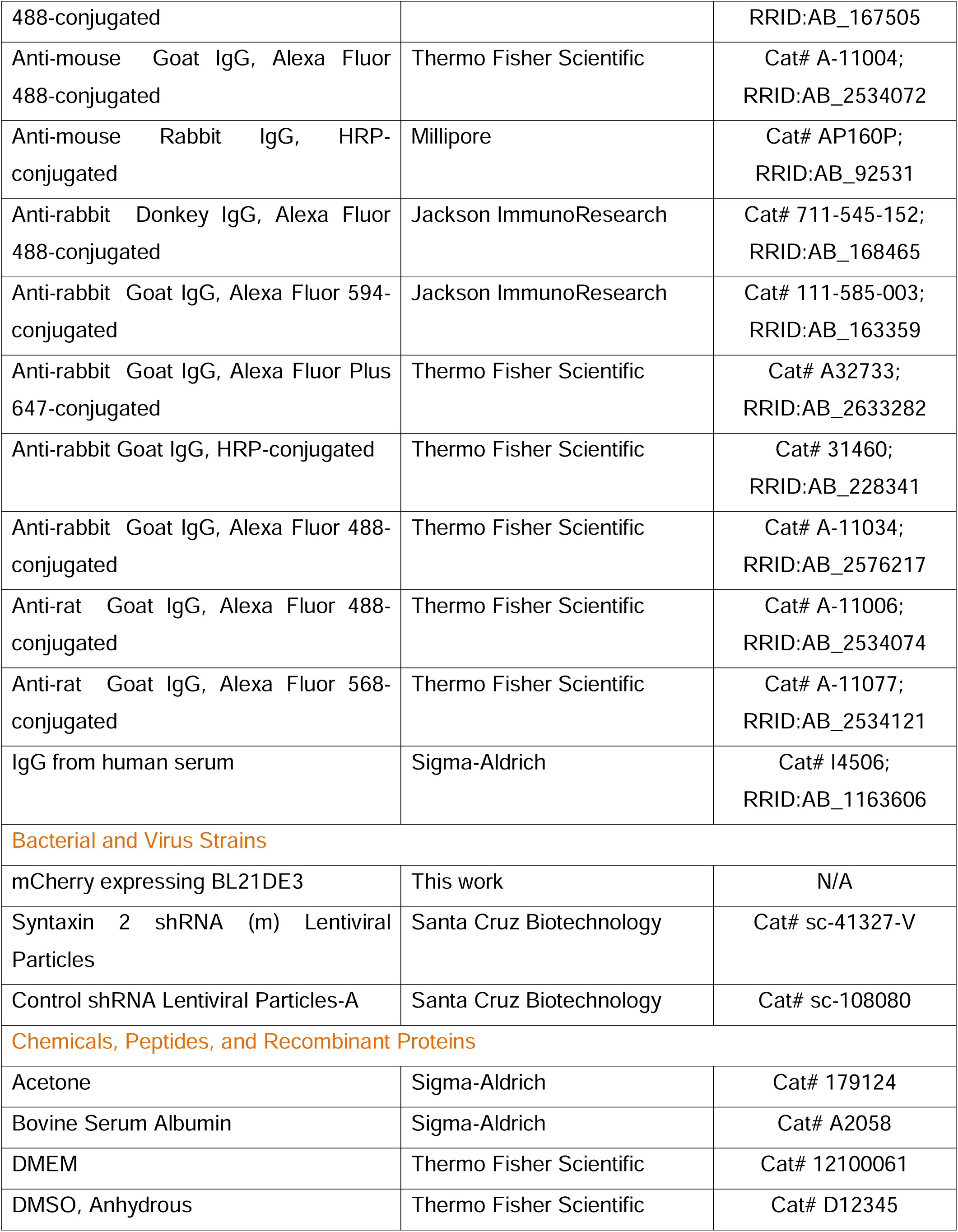
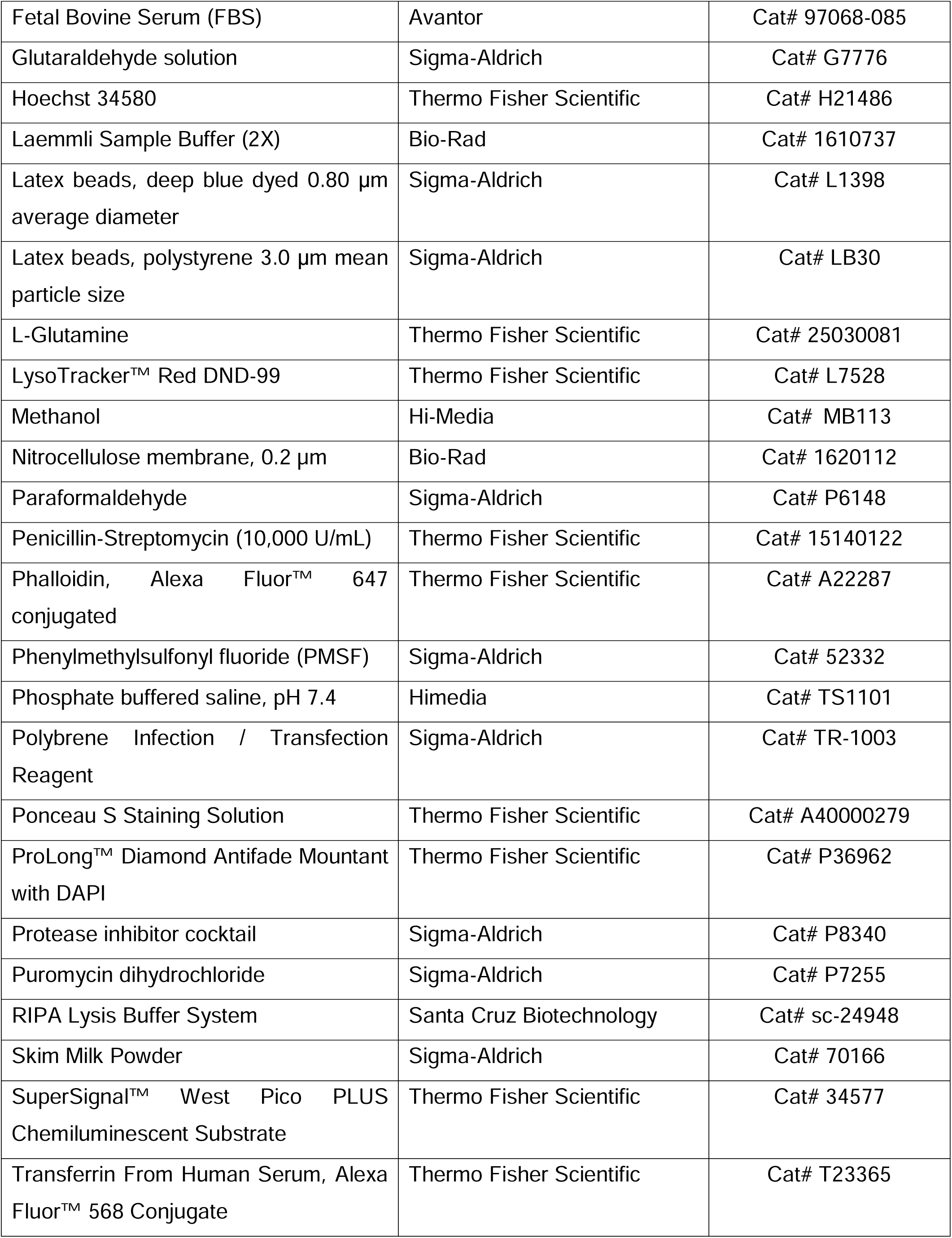
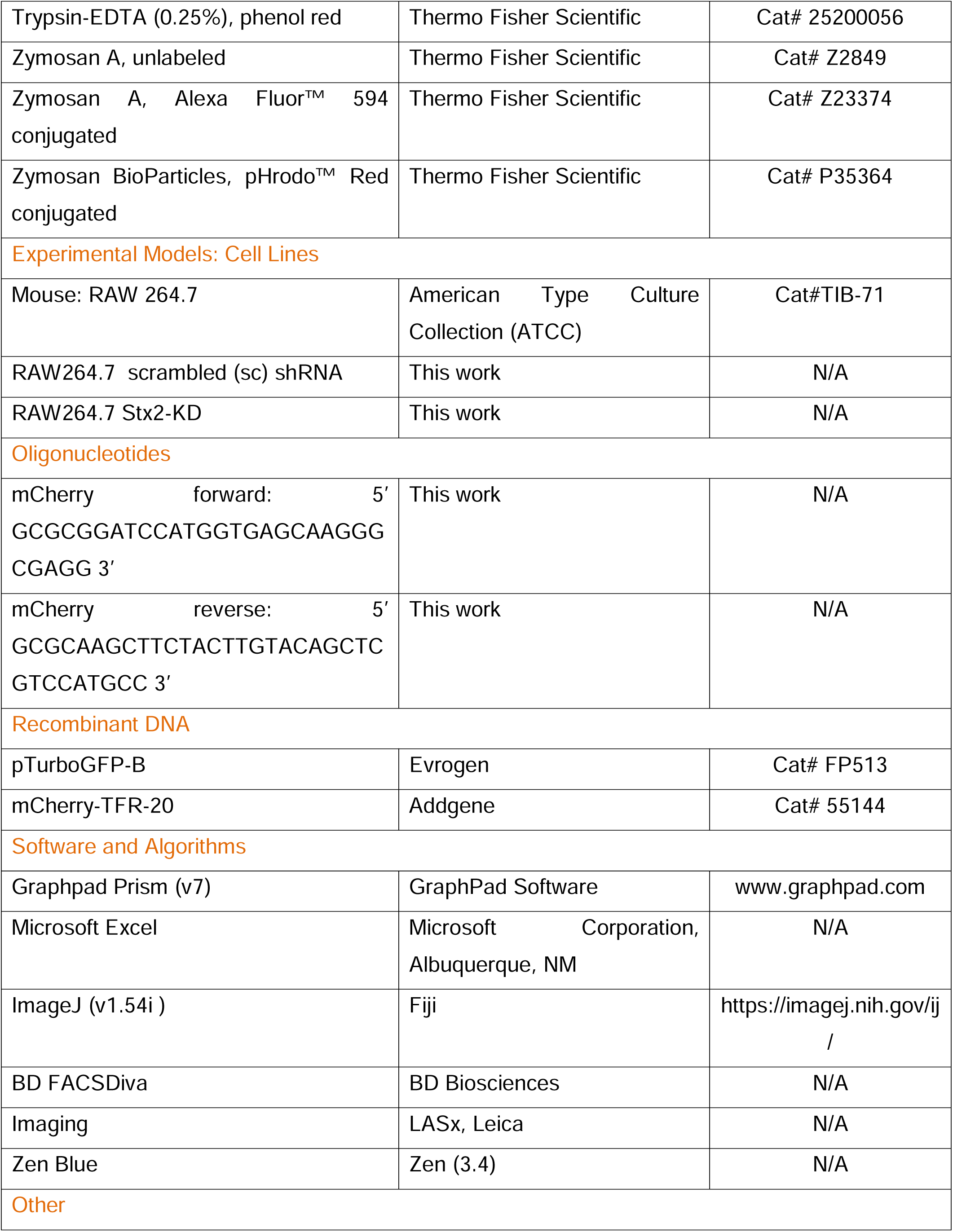
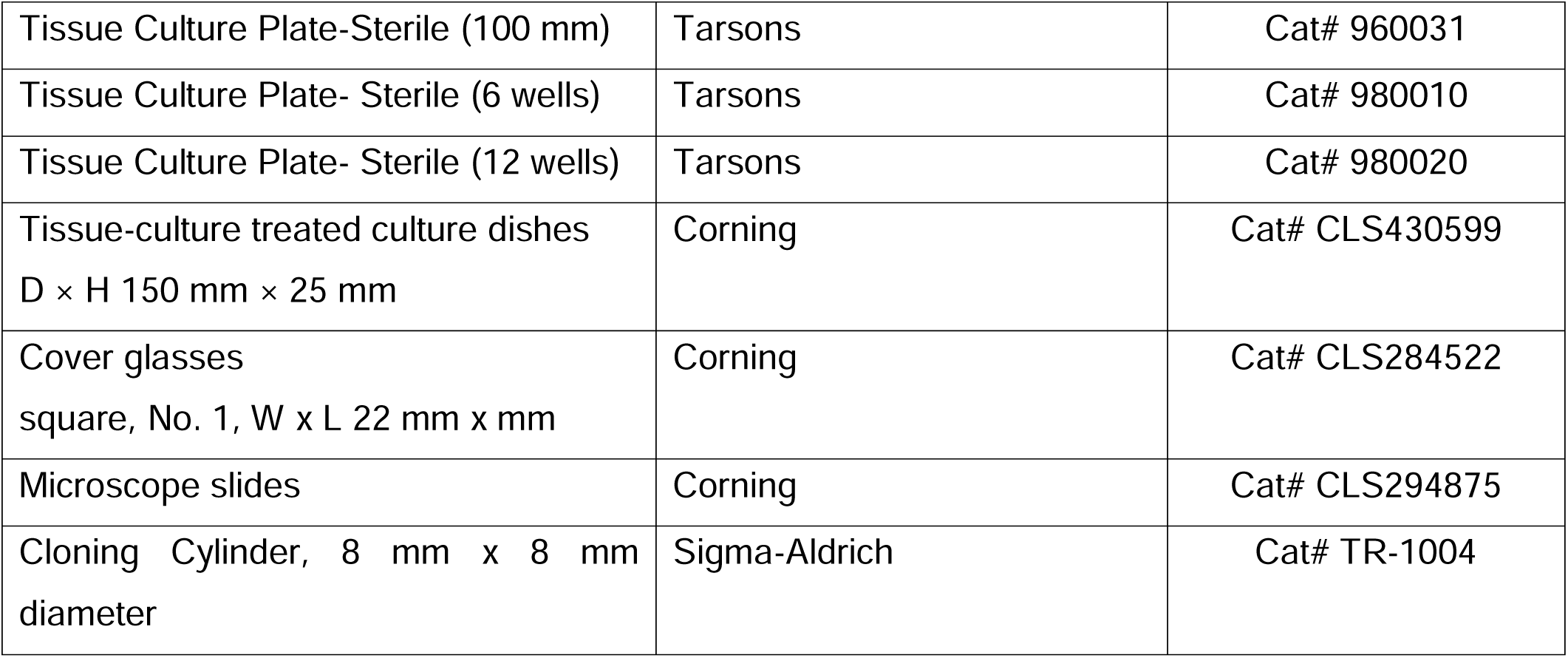
KEY RESOURCES TABLE

## RESOURCE AVAILABILITY

### Lead Contact

Further information and requests for resources and reagents should be directed to and will be fulfilled by the lead contact, Subhankar Dolai.

### Materials Availability

All the cell lines and plasmids generated for this study are available from the lead contact upon request.

### Data and Code Availability

All data reported in this paper will be shared by the lead contact upon request. This paper does not report original code. Any additional information required to reanalyze the data reported in this paper is available from the lead contact upon request.

## EXPERIMENTAL MODEL AND SUBJECT DETAILS

### Macrophage cell culture

Authentic murine RAW 264.7 macrophage cells (ATCC, catalog #TIB-71) were grown in a humidified CO_2_ (5%) incubator at 37°C in complete Dulbecco’s Modified Eagle’s Medium (DMEM; Thermo Fisher Scientific, catalog #12100061), i.e. DMEM supplemented with 10% heat-inactivated fetal bovine serum (FBS; Avantor, catalog #97068-085), 100 U/ml of penicillin-streptomycin, and 2 mM L-glutamine.

### Generation of stable Stx2 knockdown and scrambled shRNA expressing macrophages

Overnight cultures of passage 5 RAW 264.7 macrophage cells at approximately 50% confluency were mixed with 5 µg/mL polybrene (Sigma-Aldrich, catalog #TR-1003) 1 h prior to infection with lentiviral particles (MOI: 5) expressing either non-specific scrambled (sc) shRNA (Santa Cruz Biotechnology, catalog #sc-108080) to obtain Control cells or expressing mouse Stx2-shRNAs (Santa Cruz Biotechnology, catalog #sc-41327-V) to obtain Syntaxin-2 knockdown cells (Stx2- KD), following the manufacturer’s guidelines. Virus/polybrene-containing media were replaced with fresh complete DMEM 12 h after infection, and cells were allowed to grow for another 24 h before seeding onto 100 mm dishes. Puromycin (Sigma-Aldrich, catalog #P7255) was added to the culture medium at 5 µg/mL to select the transduced colonies, which were isolated individually using cloning cylinders (Sigma-Aldrich, catalog #Tr1004) and maintained under selection pressure. The expression level of Stx2 was verified by western blot, and the chosen Control and Stx2-KD clones were propagated, frozen in liquid nitrogen, and considered as a new “Passage 1”. All experiments were conducted using cells below 10 passages to minimize potential alterations due to prolonged culturing.

## METHOD DETAILS

### Generation of mCherry expressing *Escherichia coli*

The mCherry sequence was amplified from mCherry-TFR-20 plasmid (Addgene, catalog #55144) using forward primer: 5’-GCGC<u>GGATCC</u>ATGGTGAGCAAGGGCGAGG-3’, carrying BamHI restriction site (underlined) and reverse primer: 5’-GCGC<u>AAGCTT</u>CTACTTGTACAGCTCGTCCATGCC-3’, carrying HindIII restriction site (underlined). Amplified mCherry sequence was cloned into pTurboGFP-B vector (Evrogen, catalog #FP513) to replace TurboGFP reporter. Plasmids carrying mCherry were amplified and transfected into *Escherichia coli* (BL21-DE3) to produce constitutively mCherry expressing *E. coli* strain (*E. coli*-mCherry). *E. coli*-mCherry was used for phagocytosis and bacterial degradation assays.

### Phagocytosis - Preparation of Control and Stx2-KD cells

Control and Stx2-KD macrophage cells (1×10^6^ each) were seeded onto 22-mm glass coverslips (Corning, catalog #CLS284522) placed within 6-well culture plates (Tarsons, catalog #980010) containing 2 mL of 5 µg/mL puromycin (Sigma-Aldrich, catalog #P7255) mixed complete DMEM per well. Plates were kept at 37^0^C for overnight within a humidified CO_2_ (5%) incubator. Cells were washed twice (2 mL x 2) with warm (37^0^C) complete DMEM, 2 h before the initiation of phagocytosis, and finally maintained in 2 mL of warm (37^0^C) complete DMEM within a CO_2_ incubator until used.

### Phagocytosis – Preparation of opsonized phagocytic particles

Zymosan particles and latex/polystyrene beads were opsonized with human IgG (Sigma-Aldrich, catalog #I4506) 12 h before use. Unlabeled zymosan particles (Thermo Fisher Scientific, catalog #Z2849) and zymosan particles labeled with Alexa-594 (zymosan red; Thermo Fisher Scientific, catalog #Z23374) were suspended in 500 µL PBS and sonicated briefly to dissociate any aggregates. Particles were finally reconstituted in PBS (10^8^/mL) and opsonized overnight at 4^0^C by incubating with 1 mg/mL human IgG. Zymosans labeled with pHrodo red (pHRZ; Thermo Fisher Scientific, catalog #P35364) were suspended in 500 µL PBS and vortexed to dissociate any clumped particles. Reconstituted pHRZ in PBS (10^8^/mL) were supplemented with 1 mg/mL human IgG and incubated overnight at 4^0^C for opsonization. Latex/polystyrene beads of 0.8 µm diameter (Sigma-Aldrich, catalog #L1398) and 3 µm diameter (Sigma-Aldrich, catalog #LB30) were washed three times with PBS by centrifugation (5000 rpm, 5 min) and finally suspended in PBS (5×10^8^/mL) to opsonize with 1 mg/mL human IgG for overnight at 4°C. After opsonization, zymosan particles and latex/polystyrene beads were washed three times (5000 rpm, 5 min) with ice-cold PBS and stored at 4^0^C in complete DMEM. Unlabeled opsonized zymosan particles, opsonized zymosan red particles, and opsonized pHRZ were denoted as OPZ, OPZR and OpHRZ respectively. Opsonized beads of 0.8 µm and 3 µm diameters were named OPB-0.8 and OPB-3.0 respectively. *E. coli-*mCherry were opsonized with anti-LPS antibody (Cloud-Clone Corp, catalog #MAB526Ge24) 1 h before use. *E. coli*-mCherry were put into culture in Luria broth day before its usage and grown overnight at 37^0^C to reach mid-log phase (OD600 0.5-1.0). 5 mL culture was washed twice in PBS (2,000 × g, 5 min) and subjected to another round of low speed centrifugation (300 × g, 5 min) to remove any bacterial clusters. 100 µL of bacterial suspension was then opsonized with 10 µL of anti-LPS antibody for 1 h in room temperature followed by two PBS wash (2,000 × g, 5 min) and resuspension and storage at 4^0^C in complete DMEM. Opsonized *E. coli*-mCherry was abbreviated as OPEC. All opsonized particles were vortexed and quantified before use.

### Phagocytosis – Macrophage challenge with opsonized particles

Phagocytosis were performed according to published methodologies with minor adaptations^23,57^. Briefly, Control or Stx2-KD RAW 264.7 macrophages (1×10^6^ each) were seeded overnight on 22-mm glass coverslips (Corning, catalog #CLS284522) placed within 6-well culture plates (Tarsons, catalog #980010). Cells were then challenged with OPZ, OPZR, OpHRZ or OPB-3 at 15:1 (particle: cell) ratio. OPB-0.8 (intended for phagosome purification) and OPEC (intended to assay bacterial uptake and degradation) were used at 50:1(particle: cell) ratio. After addition of relevant opsonized particles, plates were immediately centrifuged at 1,000 rpm for 1 min (Hermle Z366K fitted with 221.16 rotor) to settle the particles uniformly on the macrophages. Plates were then quickly placed in a warm (37^0^C) humidified CO_2_ (5%) incubator to initiate synchronized phagocytosis. Phagocytosis events were terminated at different time points (as indicated in the Figure Legends) by replacing the media with ice-cold PBS. Particle engagement, phagocytic uptake and phagosome maturation were analyzed by fluorescence imaging, flow cytometry, phagosome fractionation, and scanning electron microscopy (SEM), where applicable, as stated below.

### Phagocytosis – Macrophage engaged particle quantification

To compare engagement of opsonized particles between Control and Stx2-KD macrophages, OPZR or OPEC challenged macrophages were washed thoroughly with ice-cold PBS at the indicated time points. Cells were then fixed with 2% PFA for 10 min followed by three PBS washes. Excess PBS was adsorbed by placing tissue papers at the edge of the coverslips. Coverslips were subsequent mounted on glass slides using a DAPI (nuclear stain) containing mounting media (Thermo Fisher Scientific, catalog #P36962). Images were acquired using a Leica SP8 confocal microscope with 63 X/1.4 NA oil immersion objectives. Both DIC and fluorescent images were recorded choosing arbitrary fields. The nuclei/cells and the fluorescent particles were counted manually and the number of engaged particles/cell were calculated by dividing total numbers of cell associated particles divided by the total numbers of nuclei. The extent of opsonized particle engagement in larger population of cells was analyzed by flow cytometry.

### Phagocytosis – Phagocytosis efficiency determination

To analyze phagocytosis efficiency, macrophage engaged particles were first distinguished as internalized and non-internalized particles by differential inside-outside staining^58^. Opsonized particle-engaged macrophages were fixed with 2% PFA for 10 min, washed thrice with PBS and blocked with 0.2% gelatin in PBS for 1h, followed by 2 h incubation with Alexa Fluor 488-conjugated anti-human secondary antibodies (1:100 dilution; for OPZR-engaged cells) or Alexa Fluor 488-conjugated anti-mouse secondary antibodies (1:100 dilution; for OPEC engaged cells), diluted in 0.2% gelatin in PBS. Secondary antibodies cannot access completely internalized OPZR or OPEC and thus can only bind to non-internalized particles to produce green/yellow fluorescence, while completely internalized particles produce only red fluorescence. Phagocytosis efficiency^58^ was calculated for each cell by using the following formula. Phagocytic efficiency _=_ Total number of internalized particles/Total number of engaged particles _*_ 100.

### Phagocytosis – Bacterial degradation assay

For bacterial degradation assay, macrophages grown on coverslips were exposed to OPEC for 2 h. Phagocytosis of OPEC was halted by ice-cold PBS wash. Macrophages were fixed and blocked as stated above, and external *E. coli* were labeled with Alexa Fluor 488-conjugated anti-mouse secondary antibodies (1:100 dilutions in 0.2% gelatin in PBS). Coverslips were mounted on glass slides and were imaged using a confocal fluorescence microscope to assess the number of engulfed bacteria (red only) and their morphology.

### Immunostaining and confocal microscopy

For immunostaining, macrophages grown overnight on coverslips were washed in PBS and fixed in 2% PFA in PBS for 10 min at room temperature (RT). Following three PBS washes (5 mins each), cells were permeabilized with 0.1% Triton X-100 in PBS for 2 min, washed thrice in PBS (5 mins each) and subsequently blocked with 0.2% gelatin in PBS (blocking buffer) for 10 min at RT. Cells intended to keep non-permeable for surface staining were skipped for permeabilization and directly used for blocking after fixation and PBS wash. Primary antibodies diluted in blocking buffer were used for 1 h to immunostain cells for Stx2 (Synaptic Systems, catalog #110123, 1:200 dilution; R and D Systems, catalog #AF2568, 1:200 dilution), Rab5A (Cell Signaling Technology, catalog #46449, 1:100 dilution), Rab11 (Cell Signaling Technology, catalog #46449, 1:200 dilution), VAMP4 (Thermo Fisher Scientific, catalog #PA1-768, 1:100 dilution), VAMP7 (Thermo Fisher Scientific, catalog #PA5-116892, 1:100 dilution), LAMP1 (Thermo Fisher Scientific, catalog #14-1071-81, 1:500 dilution), Fcγ (Thermo Fisher Scientific, catalog #14-0161-82, 1: 100 dilution), TfR (Abcam, catalog #ab214039, 1:200 dilution), Cat L (R and D Systems, catalog #AF1515, 1:100 dilution), Cat B (Cell Signaling Technology, catalog #31718, 1:100 dilution), Cat D (Proteintech, catalog #21327-1-AP, 1:100 dilution) and Phalloidin647 (Thermo Fisher Scientific, catalog # A22287, 1:500 dilution). Cells were then washed thrice in PBS (5 min each), followed by 1 hr incubation with appropriate secondary antibodies (1:800 dilution) conjugated to Alexa Fluor 488, Alexa Fluor 594 or Alexa Fluor 647. After three PBS washes, coverslips were tapped gently on tissue papers to remove excess PBS and were mounted using a mounting medium containing nuclear stain DAPI (Thermo Fisher Scientific, catalog #P36962). Z-series confocal images were captured at a 0.34 µm interval with a Leica SP8 confocal platform equipped with a 63X/1.4 NA oil immersion objective (Zeiss) with appropriate laser excitation and emission filters. Images were deconvoluted using Leica Lightning software. Fluorescence signal were quantified using Zen (Carl Zeiss) software.

### Flow cytometry (FACS)

All FACS analyses were performed by a BD LSR Fortessa flow cytometer using appropriate filter set up and analyzed with BD FACSDiva software. Lysotracker staining and FACS analyses were performed following a published methodology ^59^ with minor adjustments. Control and Stx2-KD cells were trypsinized and brought in to suspension (1.2 x 10^6^ cells/mL) in complete DMEM followed by 15 min incubation with 50 nM Lysotracker Red DND-99 at 37^0^C in a humidified CO_2_ (5%) incubator. Cells were then pelleted at 1,000 x g for 3 min, resuspended in 50 nM Lysotracker Red DND-99 containing ice-cold PBS and analyzed immediately. For surface staining of FcRs, 1.2 x 10^6^ cells were fixed in 1% PFA (in PBS) for 5 min followed by 0.2% gelatin blocking for 15 min at RT. Cells were then incubated with rat anti-FcRs/CD11b/c (1:200 dilution; Thermo Fisher Scientific, catalog #14-0161-82) antibody ^23^ for 1 h at 4^0^C followed by three PBS wash (5 min x 3) and subsequent incubation with anti-rat Alexa-488 (1:800 dilution) for 1 h at 4^0^C. Cell were again washed 3 times with PBS (5 min x 3) and finally resuspended in 1 mL PBS before FACS analyses. For the detection of phagocytic particle engagement, RAW 264.7 cells were washed with ice-cold PBS to halt uptake of fluorescent particles (OPZR and OPEC). Cells were then trypsinized and brought in to suspension in ice-cold PBS and spun down at 300 x g for 3 min at 4^0^C to remove aggregates. 0.8 mL of supernatants in ice-cold PBS containing 10^6^ cells /mL were subjected to FACS analyses.

### SEM

Monolayers of RAW 264.7 cells on glass coverslips were incubated with OPB (3 µm; 1: 15, cell: OPB) or OPEC (1: 100, cell: bacteria) and arrested for phagocytosis at different time as elaborated earlier. Cells were then fixed in 2.5% glutaraldehyde (in PBS, pH 7.4) for 1 hr at 4^0^C. After washing with PBS twice, cells were osmicated for 20 min in 1% (vol/vol) aqueous osmium tetroxide at RT. Cells were then washed with PBS, followed by dH_2_O and dehydrated by 5 min sequential incubations in 30, 50, 60, 80 and 100% ethanol. Samples were further dried overnight in a vacuum desiccator and imaged using a Zeiss Supra 55VP scanning electron microscope operating at 5.2 kV.

### Purification of phagosomes

Phagosomes were purified as described earlier with minor adjustments^43,57,60^. Briefly, Control or Stx2-KD cells were grown on 150 mm Petri dishes (Corning, catalog #CLS430599) to obtain ∼ 4×10^7^ cells for each condition. Cells were then incubated with OPB (0.8 µm, blue-dyed: OPB-0.8) at a ratio of 50:1 (beads: cell) for 30 min to 1 h. Subsequently, the cells were washed with warm DMEM to eliminate unbound OPB-0.8 and either approached for purification immediately (for 30 min OPB-0.8 incubated cells to obtain nascent phagosomes) or allowed to mature for another 30 min (for 1 h OPB-0.8 incubated cells). For phagosome isolation, cells were then washed and scrapped in ice-cold PBS, and pelleted at 1200 rpm at 4°C. Pellets were washed with 10 mL ice-cold homogenization buffer (HB: 8.55% sucrose, 3 mM imidazole, 2X PIC, pH 7.4) for 5 mins and pelleted at 1200 rpm. Pellets were brought to suspension in 1 mL HB with 2X PIC and passed through a 30-G needle (fitted in a 1 mL syringe) until L90% of cells were broken, as confirmed by phase microscopy. The homogenates were clarified at 1200 rpm for 5 mins and supernatants enriched with OPB-0.8 containing phagosomes were mixed with an equal volume of 62% sucrose. This mixture was layered over 3 mL of 62% sucrose, followed by sequential topping with 2 mL each of 35, 25, and 10% sucrose (all prepared in 3 mM imidazole, pH 7.4) in SW40 tubes (344060, Beckman). The samples were then centrifuged at 100,000 x g for 1 h at 4°C using a SW40Ti rotor (Beckman). Phagosomes were recovered from the interface of 10 and 25% sucrose (Fig. S4) using a pipette, washed with 12 mL ice-cold PBS, and harvested at 40,000 x g.

### Subcellular fractionation

∼ 90% confluent RAW 264.7 cells in 100 mm dishes were washed twice in PBS and scrapped into 200 µL of NP-40 lysis buffer (l0 mM Tris, pH 7.9, 140 mM KCl, 5 mM MgCl2 and 0.5 % NP-40)^61^ supplemented with PIC and kept on ice for 15 min. Cells were passed through a 22G syringe for 5 to 10 times until ≤90% of cells were broken (verified under the light microscope). Lysates were then spun at 100 x g for 5 min to pellet down unbroken cells. Supernatants were collected and further centrifuged at 1000 x g for 5 min to pellet down nucleus. The resultant supernatant represents the cytosolic plus the membrane fraction. The nuclear pellets were washed twice in NP-40 lysis buffer and then sonicated in Laemmli sample buffer equal to the volume of cytosolic/membrane fraction. Equal volume of nuclear and cytosolic/membrane fractions were clarified by SDS-PAGE along with 20 µg of TCL.

### Cathepsin secretion assay

Equal number (1 X 10^7^) of Control and Stx2-KD RAW 264.7 macrophages were seeded on 100 mm tissue culture plates and allowed to adhere for 6 h. Macrophages were then washed three times with serum-free DMEM to remove any non-adherent cells and finally kept in 5 mL of serum-free DMEM for 12 h. Conditioned media were collected and subjected to centrifugation (1000 x g, 5 min) to remove any cells. 4 mL of each conditioned medium were then subjected to acetone (Sigma-Aldrich, catalog #179124) precipitation by mixing with 20 mL of pre-cooled (−20 C) acetone, followed by 2 h incubation at -20 C. Precipitated proteins were pelleted at 13,000 × g for 20 min at 4°C. Pellets were air-dried to remove traces of acetone and finally dissolved in 200 µL of Laemmli buffer. 20 µL of each sample were clarified by SDS-PAGE along with 20 µg of total cell lysates. Presence of cathepsins in secretion was detected by western blot.

### Western blotting

RAW 264.7 cells or purified phagosomes were lysed in ice-cold RIPA buffer (Santa Cruz Biotechnology; catalog #sc-24948) with a protease inhibitor cocktail (PIC, 1X; Sigma-Aldrich, catalog #P8340). Samples were then sonicated (Sartorius) and separated from insoluble debris at 10,000g for 10 min at 4°C. Protein concentration was estimated using modified Lauri method and brought to similar level by adding RIPA buffer containing PIC. Lysates were then boiled in Laemmli buffer (1.5X), resolved by 10-12% SDS–PAGE and transferred to nitrocellulose membranes using a semi-dry transfer system (BioRad). Non-specific binding sites were blocked in 5% skim milk dissolved in TBST (TBS: 50 mM Tris-HCl [pH 7.4], 274 mM NaCl, 9 mM KCl with 0.05% Tween-20) for 1 h at RT and membranes were washed briefly to remove excess milk. Primary antibodies diluted in 1^0^ antibody buffer (1% BSA [w/v] in 50 mM Tris-HCl [pH 7.4]) were used to probe membranes for Stx2 (Thermo Fisher Scientific, catalog #PA5-87903, 1:1000 dilution), Rab5A (Cell Signaling Technology, catalog #46449; 1:1000 dilution), VAMP3 (Proteintech, catalog #10702-1-AP, 1:1000 dilution), VAMP4 (Thermo Fisher Scientific, catalog #PA1-768, 1:1000 dilution), LAMP1 (R and D Systems, catalog #AF4320, 1:1000 dilution), Golgin 160 (Santa Cruz Biotechnology, catalog #sc-374596, 1:500 dilution), VAMP7 (Thermo Fisher Scientific, catalog #PA5-116892, 1:1000 dilution), VAMP8 (Proteintech, catalog #15546-1-AP, 1:1000 dilution), Cathepsin L (R and D Systems, catalog #1515, 1:1000 dilution), Cathepsin B (Cell Signaling Technology, catalog #31718, 1:1000 dilution), Cathepsin D (Proteintech, catalog #21327-1-AP, 1:1000 dilution), ATP6V1A (Cell Signaling Technology, catalog #39517, 1:1000 dilution), TFEB (Thermo Fisher Scientific, catalog #PA5-96632, 1:1000 dilution), Histone H3 (Cell Signaling Technology, catalog #9715, 1:2000 dilution), LC3A/B (Cell Signaling Technology, catalog #12741, 1:400 dilution), SQSTM1/p62 (Cell Signaling Technology, catalog #39749, 1:800 dilution) β-actin (Cell Signaling Technology, catalog #3700, 1:1000 dilution) and Tubulin (Cell Signaling Technology, catalog #3873; 1:1000 dilution) for 12 h at 4^0^C. After washing three times (10 min X 3) with TBST, membranes were incubated with suitable HRP-conjugated secondary antibodies diluted (1: 5000) in 5% skim milk/TBST for 1 h at RT. Membranes were then washed three times in TBST (10 min X 3), developed with ECL reagent and documented using a ChemiDoc imaging system (Bio-Rad). Membranes containing phagosome samples were stained with Ponceau S (5 min) after semi-dry transfer to document loading and used to wash with TBST to remove stain before proceeding for blocking. Densitometry analyses of the western blot bands were performed using ImageJ 1.54i software (NIH, USA). Band density values were first normalized to loading controls and values from Controls were then adjusted to 1. All other values were expressed as relative to Control values by dividing with 1.

## QUANTIFICATION AND STATISTICAL ANALYSIS

### Measuring phagocytic cup area

Acquired SEM images were opened in ImageJ, and the scale was calibrated to ensure area calculation in micrometers and that the phagocytic cup region in each macrophage was manually outlined using the freehand selection tool in ImageJ. Then the area of the selected region was measured in µm² using the “measure” command.

### IF intensity quantification

Fluorescence intensity (in A.U.) for each data points in Control cells were normalized by dividing with average value. This normalization step transformed the control values into relative values centered on 1. Average value from the Control was then used to normalize the raw intensity values (in A.U.) from the treatment groups by dividing each raw value by the Average Control value.

### Western blot densitometry

Band density values from control samples were normalized to 1 and the band density values from treated samples were divided by 1 to generate relative density values.

### Statistics

Two-tailed Student’s t-test statistical analyses were performed using GraphPad Prism software (v.9.3.0). All the values are expressed as means ± SEM. All experiments were repeated independently for at least three times. The actual numbers of experiments and cells or phagosomes quantified are mentioned in the individual Figure Legends.

## SUPPLEMENTAL FIGURE LEGENDS

**Figure S1. Lenti/Stx2-shRNAs significantly deplete endogenous Stx2 in RAW264.7 macrophages and increase macrophage capacity for IgG-opsonized particle engagement.** (A) Representative western blots from Lenti/sc-shRNA transduced (Control) or Lenti/Stx2-shRNA transduced (Stx2-KD) RAW 264.7 cells to test Stx2 depletion. C1-C3 and C5-C7 represents the parent puromycin resistant colonies. Tubulin used as loading control. (B) Quantification of Stx2 band densities normalized to tubulin. “*” indicates the colony 1 (C1) with maximum Stx2 depletion (Stx2-KD: ∼67%) that further amplified and used in all the experiments stated in this article. (C) Representative DIC (upper panel) and fluorescence images (bottom panel) created by stacking 3 consecutive optical sections from Control or Stx2-KD macrophages challenged with IgG-opsonized fluorescent beads (OPFB, 2 µm diameter: OPFB-2) for 15, 30 and 60 min. DAPI (blue) shows the nuclei. Scale bars, 10 μm. (D) Quantification of the macrophage engaged OPFB-2 in Control and Stx2-KD macrophages (15min Control, n=20; Stx2-KD, n= 21),(30min Control, n=25 ; Stx2-KD, n= 20), (60min Control, n=25 ; Stx2-KD, n= 27) N = 3 independent experiments. Error bars, ± SEM. ***P < 0.001.

**Figure S2. Increased OPB-3 engagement on Stx2-KD macrophage surface.** Additional SEM images of Control (upper panel) and Stx2-KD (bottom panel) macrophages incubated with OPB-3 for the indicated time. Stx2-KD cells are showing more efficiency for OPB attachment.

**Figure S3: Stx2 depletion augments macrophage lysosome content**. (A) Representative confocal DIC and fluorescence images of Control (top panel) and Stx2-KD (bottom panel) RAW 264.7 cells stained with LysoTracker Red DND-99. Boxed regions are magnified in inset. Nucleus (blue) stained with Hoechst. Scale bars, 10 μm, 5 µm (inset). (B) Quantification of LysoTracker Red intensity (in arbitrary unit: A.U.) from individual Control and Stx2-KD RAW 264.7 macrophages (Control, n= 93; Stx2-KD, n= 118) presented as violin plot. N = 3 independent experiments. Error bars, ± SEM. ***P < 0.001. (C) Representative western blots show expression of LAMP1 and loading control ACTB in total cell lysates (TCL) from Control and Stx2-KD macrophages. (D) Quantification of LAMP1 band density normalized to ACTB loading control. N = 3 independent experiments. Error bars, means ± SEM. **P < 0.01. (E) Representative western blots of total cell lysates (TCL) from Control and Stx2-KD RAW 264.7 macrophages probed for LC3-I/II, SQSTM1/p62 and loading control ACTB. (F and G) Quantification of band density for (F) LC3-II/I and (G) SQSTM1/p62 normalized to ACTB. N = 3 independent experiments. Error bars, means ± SEM. **P < 0.01. (H) Quantification of TFEB band densities in TCL, nucleus and cytosol. N = 3 independent experiments. Error bars, means ± SEM. ns = not significant.

**Figure S4. Schematic presentation of the phagosome purification methodology, described in detail in the “materials and method” section.** OPB-0.8 containing phagosomes were purified by using sucrose density gradient centrifugation as outlined.

