## Supplementary Data for "Syntaxin-2 coordinates endolysosomal trafficking to balance phagocytic uptake and phagolysosomal clearance in macrophages"

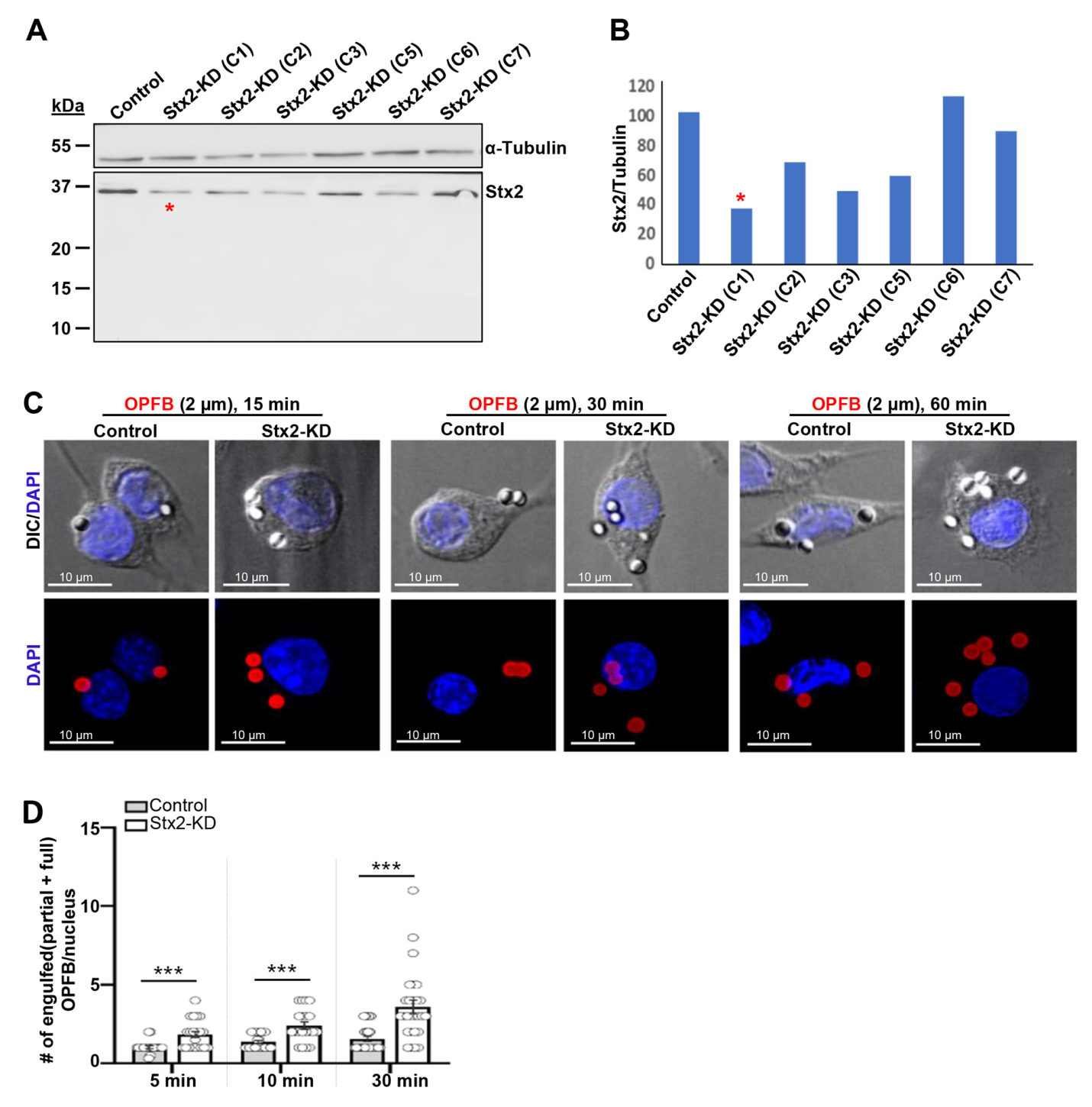

**Figure S1. Lenti/Stx2-shRNAs significantly deplete endogenous Stx2 in RAW264.7 macrophages and increase macrophage capacity for IgG-opsonized particle engagement.** (A) Representative western blots from Lenti/sc-shRNA transduced (Control) or Lenti/Stx2-shRNA transduced (Stx2-KD) RAW 264.7 cells to test Stx2 depletion. C1-C3 and C5-C7 represents the parent puromycin resistant colonies. Tubulin used as loading control. (B) Quantification of Stx2 band densities normalized to tubulin. “*” indicates the colony (C1) with maximum Stx2 depletion (Stx2-KD: ~67%) that further amplified and used in all the experiments stated in this article. (C) Representative DIC (upper panel) and fluorescence images (bottom panel) created by stacking 3 consecutive optical sections from Control or Stx2-KD macrophages challenged with IgG-opsonized fluorescent beads (OPFB, 2 µm diameter: OPFB-2) for 15, 30 and 60 min. DAPI (blue) shows the nuclei. Scale bars, 10 μm. (D) Quantification of the macrophage engaged OPFB-2 in Control and Stx2-KD macrophages (15min Control, n=20; Stx2-KD, n= 21),(30min Control, n=25 ; Stx2-KD, n= 20), (60min Control, n=25 ; Stx2-KD, n= 27) N = 3 independent experiments. Error bars, ± SEM. ***P < 0.001.

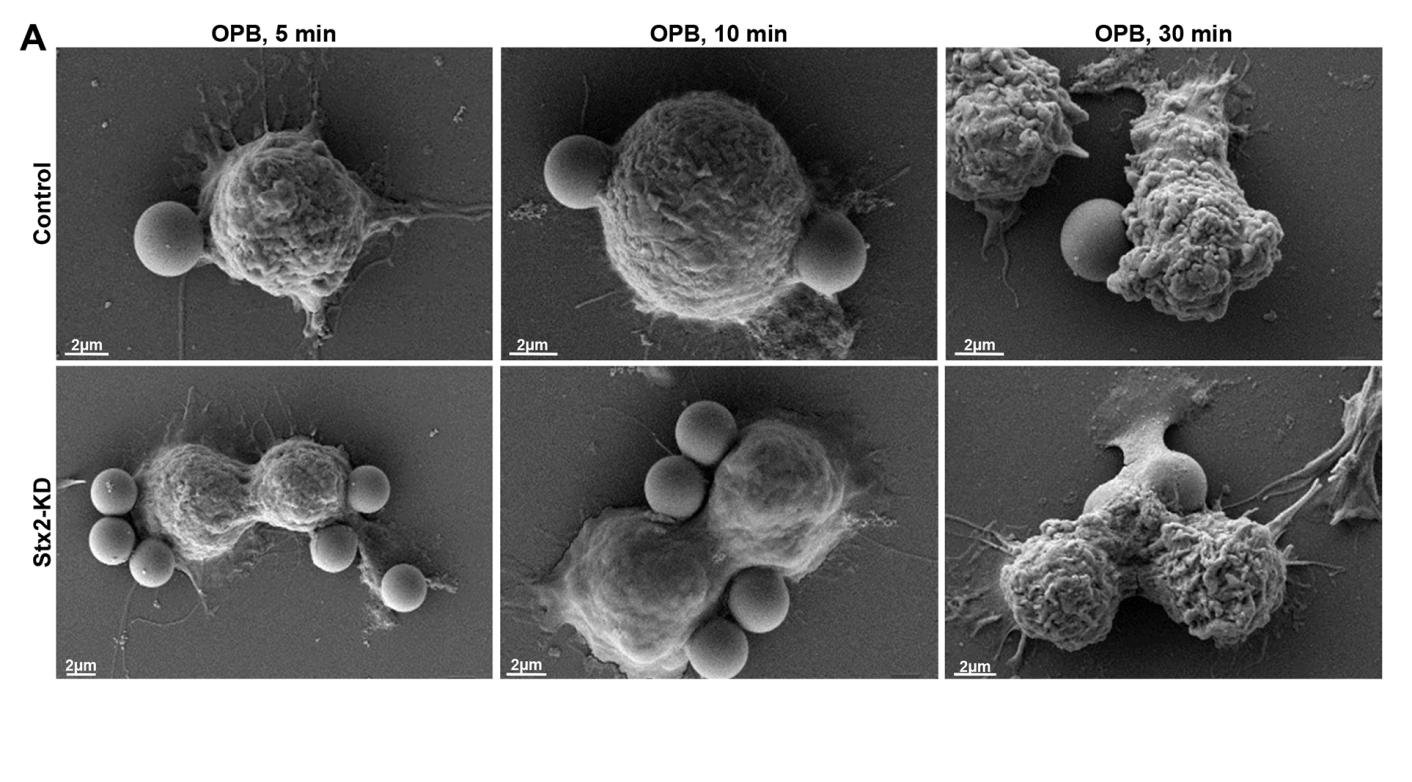

**Figure S2.** **Increased OPB-3 engagement on Stx2-KD macrophage surface.** Additional SEM images of Control (upper panel) and Stx2-KD (bottom panel) macrophages incubated with OPB-3 for the indicated time. Stx2-KD cells are showing more efficiency for OPB attachment.

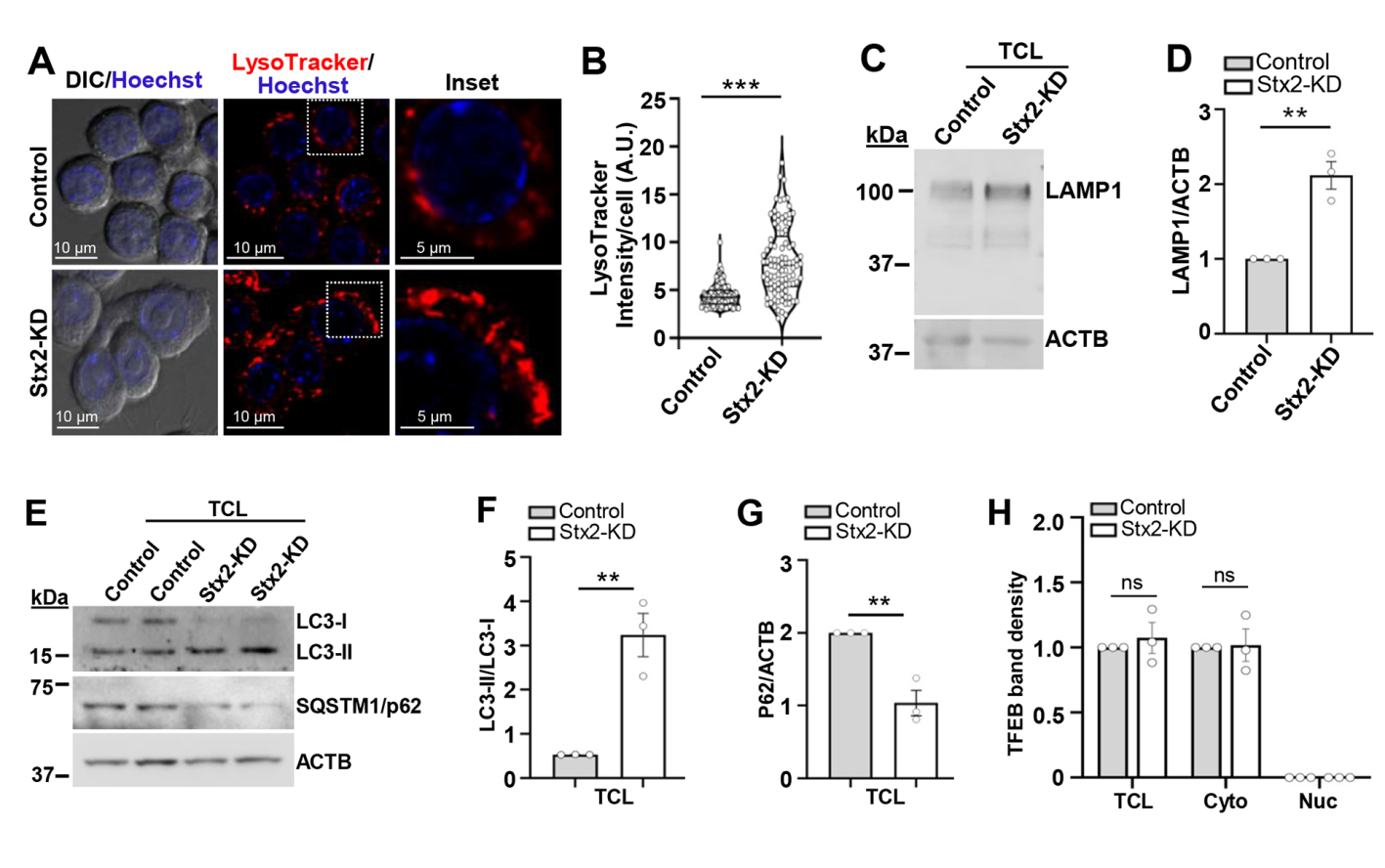

**Figure S3: Stx2 depletion augments macrophage lysosome content**. (A) Representative confocal DIC and fluorescence images of Control (top panel) and Stx2-KD (bottom panel) RAW 264.7 cells stained with LysoTracker Red DND-99. Boxed regions are magnified in inset. Nucleus (blue) stained with Hoechst. Scale bars, 10 μm, 5 µm (inset). (B) Quantification of LysoTracker Red intensity (in arbitrary unit: A.U.) from individual Control and Stx2-KD RAW 264.7 macrophages (Control, n= 93; Stx2-KD, n= 118) presented as violin plot. N = 3 independent experiments. Error bars, ± SEM. ***P < 0.001. (C) Representative western blots show expression of LAMP1 and loading control ACTB in total cell lysates (TCL) from Control and Stx2-KD macrophages. (D) Quantification of LAMP1 band density normalized to ACTB loading control. N = 3 independent experiments. Error bars, means ± SEM. **P < 0.01. (E) Representative western blots of total cell lysates (TCL) from Control and Stx2-KD RAW 264.7 macrophages probed for LC3-I/II, SQSTM1/p62 and loading control ACTB. (F and G) Quantification of band density for (F) LC3-II/I and (G) SQSTM1/p62 normalized to ACTB. N = 3 independent experiments. Error bars, means ± SEM. **P < 0.01. (H) Quantification of TFEB band densities in TCL, nucleus and cytosol. N = 3 independent experiments. Error bars, means ± SEM. ns = not significant.

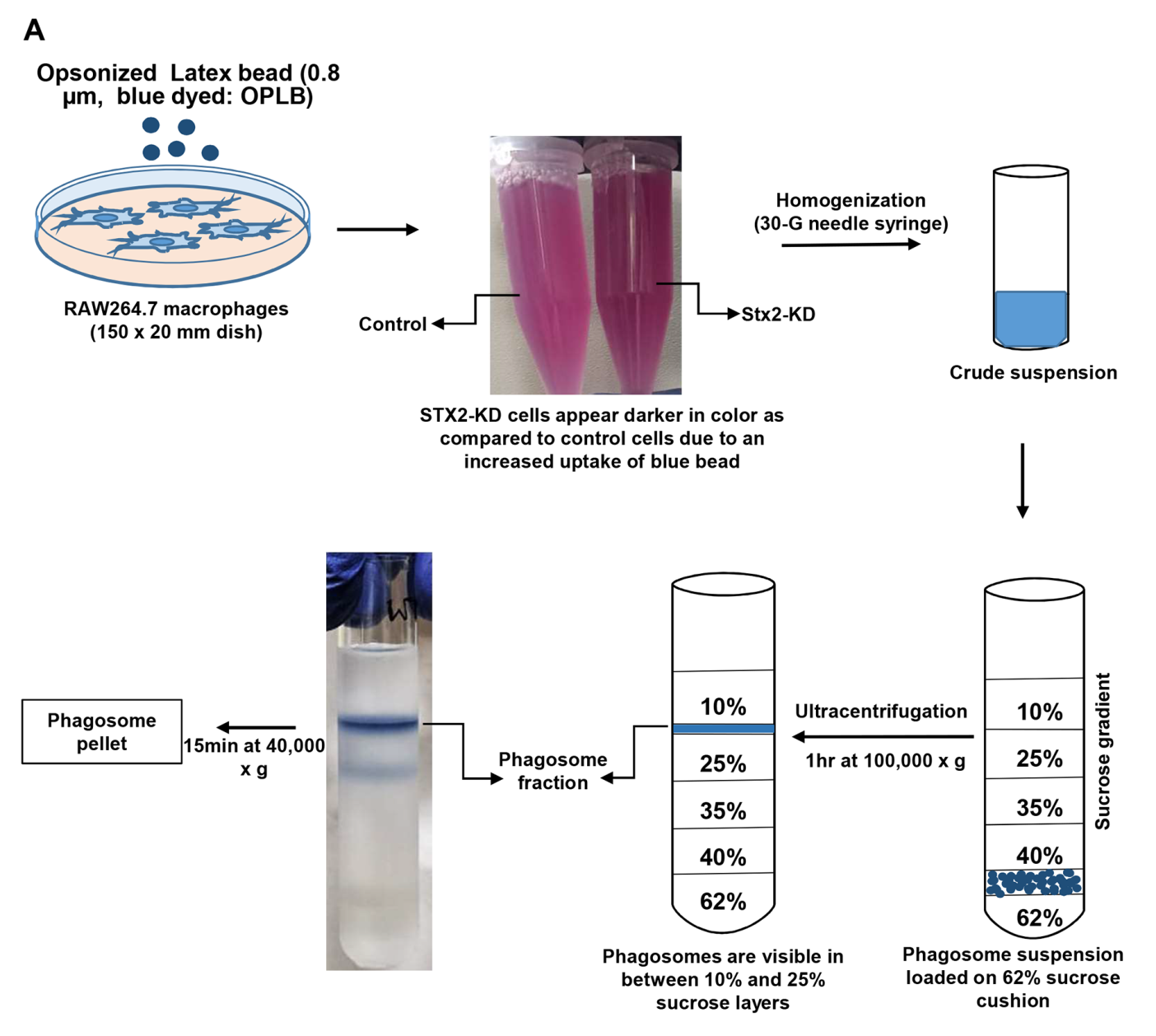

**Figure S4. Schematic presentation of the phagosome purification methodology, described in detail in the “materials and method” section.** OPB-0.8 containing phagosomes were purified by using sucrose density gradient centrifugation as outlined.
